# Host behavior alteration by its parasite: from brain gene expression to functional test

**DOI:** 10.1101/2020.05.08.084764

**Authors:** Lucie Grecias, Francois Olivier Hebert, Verônica Angelica Alves, Iain Barber, Nadia Aubin-Horth

## Abstract

Many parasites with complex life cycles modify their intermediate hosts’ behaviour, presumably to increase transmission to their final host. The threespine stickleback (*Gasterosteus aculeatus*) is an intermediate host in the cestode *Schistocephalus solidus* life cycle, which ends in an avian host, and shows increased risky behaviours when infected. We studied brain gene expression profiles of sticklebacks infected with *S*.*solidus* to determine the proximal causes of these behavioural alterations. We show that infected fish have altered expression levels in genes involved in the inositol pathway. We thus tested the functional implication of this pathway and successfully rescued normal behaviours in infected sticklebacks using lithium exposure. We also show that exposed but uninfected fish have a distinct gene expression profile from both infected fish and control individuals, allowing us to separate gene activity related to parasite exposure from consequences of a successful infection. Finally, we find that Selective Serotonin Reuptake Inhibitor (SSRI)-treated sticklebacks and infected fish do not have similarly altered gene expression, despite their comparable behaviours, suggesting that the serotonin pathway is probably not the main driver of phenotypic changes in infected sticklebacks. Taken together, our results allow us to predict that if *S*.*solidus* directly manipulates its host, it could target the inositol pathway.

## Introduction

Many parasites go through complex life cycles and as they do so alter various aspects of their host’s biology, including morphology, physiology, life history, and behaviour [1]. These phenotypic changes can decrease the host’s fitness and have been proposed to increase the probability of completion of the parasite’s life cycle [2, 3], although in many cases experimental evidence is still needed [1]. Host behaviour manipulations can range from slight changes in pre-existing traits to the display of entirely novel behaviours [4]. A striking example is provided by the threespine stickleback (*Gasterosteus aculeatus*) and its tapeworm parasite, *Schistocephalus solidus*. The parasite has three hosts: a copepod, a fish (specifically the threespine stickleback), and a fish-eating bird, the definitive host [5-7]. The presence of *S. solidus* in the body cavity of its host has been reported to have multiple effects on stickleback physiology, including increased oxygen consumption [8], reduced gonad development [9], and decreased energy reserves [10]. Infected sticklebacks lose their anti-predator response and forage under the risk of predation [11-13]. They spend less time swimming within a group than healthy conspecifics [14], tend to swim away from cover, a sign of lower anxiety [18] and tend to swim close to the surface [15] even during the day, which is rarely seen in healthy conspecifics [16].

Characterizing the mechanistic basis of the interaction between the parasite and its host in the context of behavioural change requires three steps. The first one is to uncover which molecular pathways are altered in parasitized hosts. The second step is to use experimental manipulations to single out which molecular changes are the cause of the changes in behaviour [17, 18]. After confirming the causal role of a molecular pathway in behaviour variation, the third step is to determine which one of these pathways, if any, is directly manipulated by the parasite [19]. Here we present data in the stickleback-*S. solidus* host-parasite system, obtained during experiments pertaining to the two first steps. Despite a rich literature describing altered host phenotypes, there is comparatively less information on the proximate pathways involved in drastic behavioural changes in most host-parasite pairs [20, 21], including in the stickleback-*S. solidus* system [7]. We can predict that these mechanistic bases include interconnected levels of biological organization: neural circuits, neuroendocrine regulation (potentially including the serotonergic axis [22], see below), gene expression changes, and epigenetic regulation [23]. Because of the multidimensional nature of phenotypic changes in the parasitized sticklebacks that include several types of behaviours, but also physiology [24, 25] and immunity [26, 27], an assumption-free whole-genome approach to characterize gene expression changes in the brain is optimal [28-30]. We can also expect that the host responds to infection [26, 31] and that this will be reflected in the gene expression profiles, as found in innate and adaptive immune system genes of the stickleback’s head kidney [32]. The stickleback-*S. solidus* host-parasite system is an excellent model to study the genomic signature of parasitic infection in the host brain, i.e. a group of genes with a characteristic pattern of expression that occurs as a result of a biological process [33-35]. Sticklebacks can be experimentally infected, allowing the control of other environmental variables that could affect control and infected fish [36].

Changes in gene expression of an infected host compared to a non-infected conspecific might be functionally associated with the behavioural changes observed but could also merely be the consequence of being exposed to a parasite. An important question thus arises: do individuals exposed to a parasite that did not become infected have a similar brain expression profile to control individuals, to infected hosts, or is it unique? Since not all stickleback exposed to *S. solidus* become infected [7], it is possible to also study gene expression profiles of these individuals. While it has been shown that head kidney gene expression patterns do not differ between control and exposed sticklebacks [32], there are no available studies on the brain genomic signatures of exposed individuals to test these contrasting predictions.

One approach to address the question of differences in brain genomic signatures is through the study of the host’s serotonergic neuroendocrinological pathway, which may be modified indirectly or directly by behaviour-altering parasites. Studies in various systems have shown changes in candidate molecules such as biogenic amines in parasitized individuals (insects: [37, 38], crustaceans: [39], fish: [40]). In stickleback, serotonin activity is higher in *S. solidus*-parasitized wild-caught female sticklebacks compared to healthy females, which has been attributed to the stress of being parasitized [22]. Furthermore, experimental pharmacological manipulation of biogenic amines such as the serotonergic axis in healthy individuals results in behavioural changes typical of infected host (in crustaceans, [17]). In sticklebacks, Selective Serotonin Reuptake Inhibitor (SSRI)-treated non-infected sticklebacks show similar behaviours to *S. solidus*-infected individuals, with a lower tendency to school with conspecifics, and more time spent at the surface, although only in some individuals, while anti-predator response is not affected [18]. Since behavioural changes in parasitized individuals overlap in part with the ones measured in SSRI-treated individuals, they could exhibit similar activity of certain molecular pathways, which can be quantified by comparing their brain gene expression profiles.

Here, we investigated genome-wide brain gene expression patterns of sticklebacks from four treatments using RNA-seq: healthy controls, infected by *S. solidus*, exposed to a *S. solidus* parasite but not infected, and SSRI-treated. First, we analysed the transcriptome of *S. solidus-*infected stickleback. We predicted that they would show changes in expression of genes related to the multidimensional phenotypic changes they exhibit: behaviour, physiological systems and host response to infection. We then performed a follow-up experiment using a pharmacological manipulation, to test the behavioural effects of manipulating a candidate molecule found to be highly expressed in the brain of infected sticklebacks. Second, we included exposed individuals in which worms did not develop. We predicted that exposed fish would have a gene expression pattern mostly related to the host response to infection, which would match a subset of the expression profile of a successfully infected stickleback. Finally, we used individuals treated with the SSRI fluoxetine. We predicted that if *S. solidus* affects the same molecular pathways as the SSRI, we would detect a high overlap when comparing brain gene expression profiles of SSRI-treated versus infected fish.

## Materials and Methods

### Exposure of fish host to its parasite or SSRI

Sticklebacks from Llyn Frongoch (UK) were bred and their offspring reared in the laboratory for six months (see [18] and supplementary material). In summary, we created four treatment groups. We exposed individuals to *S*.*solidus*-infected copepods (see [18] for infection techniques) and waited three months for parasite growth. This treatment resulted in two groups: infected (fish with a parasite) and exposed fish (fish without a parasite). It was not possible to distinguish the exposed and infected individuals prior to dissection. We also exposed individuals to the SSRI fluoxetine for three days at a dose of 1mg / L (Fluoxetine HCl, BML-NS140, Enzo Life Sciences Inc., USA), known to result in behavioural changes similar to those induced by the presence of *S. solidus* (see [18] for details). Control fish that were never exposed to a parasite were kept in the same conditions in parallel. Before fish were euthanized, they were screened for ecologically-relevant behaviours, but the small sample size for infected fish (n=3) prevented meaningful statistical analyses (see [18]). The parasites found in the infected sticklebacks were confirmed to be in the infective stage using their transcriptome profiles [18].

### Gene expression quantification by RNA-seq

Fish were euthanized following authorised protocol and dissected brains were kept in RNALater (Ambion Inc., Austin, TX, USA). We extracted total RNA from the brains of three infected, six exposed, six SSRI-treated, and six control fish (all females) using a standard Trizol reagent protocol (miRNeasy Micro kit, Qiagen) and stored at −80°C after verifying concentration and quality by spectrophotometer (Nanodrop, Thermo scientifics) and a Bioanalyzer (RNA 6000 Nano Kit, Agilent Technologies Inc). We produced libraries for these 21 individuals using the TruSeq RNA Library Prep Kit v2 (Illumina, Inc., USA) with a unique barcode for each library. Library quality and size was assessed on a Bioanalyzer High Sensitivity DNA Assay (Agilent Technologies). The 21 cDNA libraries were then pooled and sequenced (Illumina HiSeq 2000). See supplementary material for details.

### Analysis of differential gene expression

The complete RNA-Seq data preparation pipeline is available in details in supplementary material. We used the R packages “edgeR” 3.24.3 [41] and “limma-voom” v.3.7 [42] to filter the dataset and determine differential gene expression. After quality control and data filtering, we used 12,520 annotated transcripts and 20 of the 21 original libraries (one exposed individual was removed because of poor quality, see supplementary figure 1). Absolute read counts were converted into their respective CPM value and log2-transformed using the “voom” function. Each transcript was fitted to an independent linear model using the log2(CPM) values as the response variable and the treatment as the explanatory variable. Each linear model was then analysed through limma’s Bayes pipeline. We determined which genes were differentially expressed in each group (infected, exposed, SSRI-treated) compared to healthy controls based on a p-value of p<0.005. We did not apply a false discovery rate correction, as it greatly reduced our dataset, with the caveat that interpretation of changes in expression of a specific gene must be done only as a preliminary result and an additional functional analysis is needed to corroborate our findings (which we did for one candidate gene, see “Functional analysis” section and discussion). The results of statistical comparisons between control individuals and each treatment with associated fold-change and p-value are in supplementary tables S1, S4 and S6. Within differentially expressed genes, we identified genes that are differentially expressed only in that specific treatment vs the control group, to define a genomic signature of that treatment group (ex: significantly more expressed in infected fish compared to controls, but not differentially expressed between exposed fish and controls, or between SSRI-treated fish and controls) [29, 35]. These genes are marked in bold in the corresponding supplementary tables. We performed an enrichment analysis for each genomic signature separately, to test if certain biological functions were significantly overrepresented. GO terms for each gene were based on the published transcriptome of *Gasterosteus aculeatus*. We used the Python package ‘goatools’ v.0.6.5 [43] to perform Fisher’s exact tests using a p-value of p<0.005 as a significance threshold.

**Figure 1.**
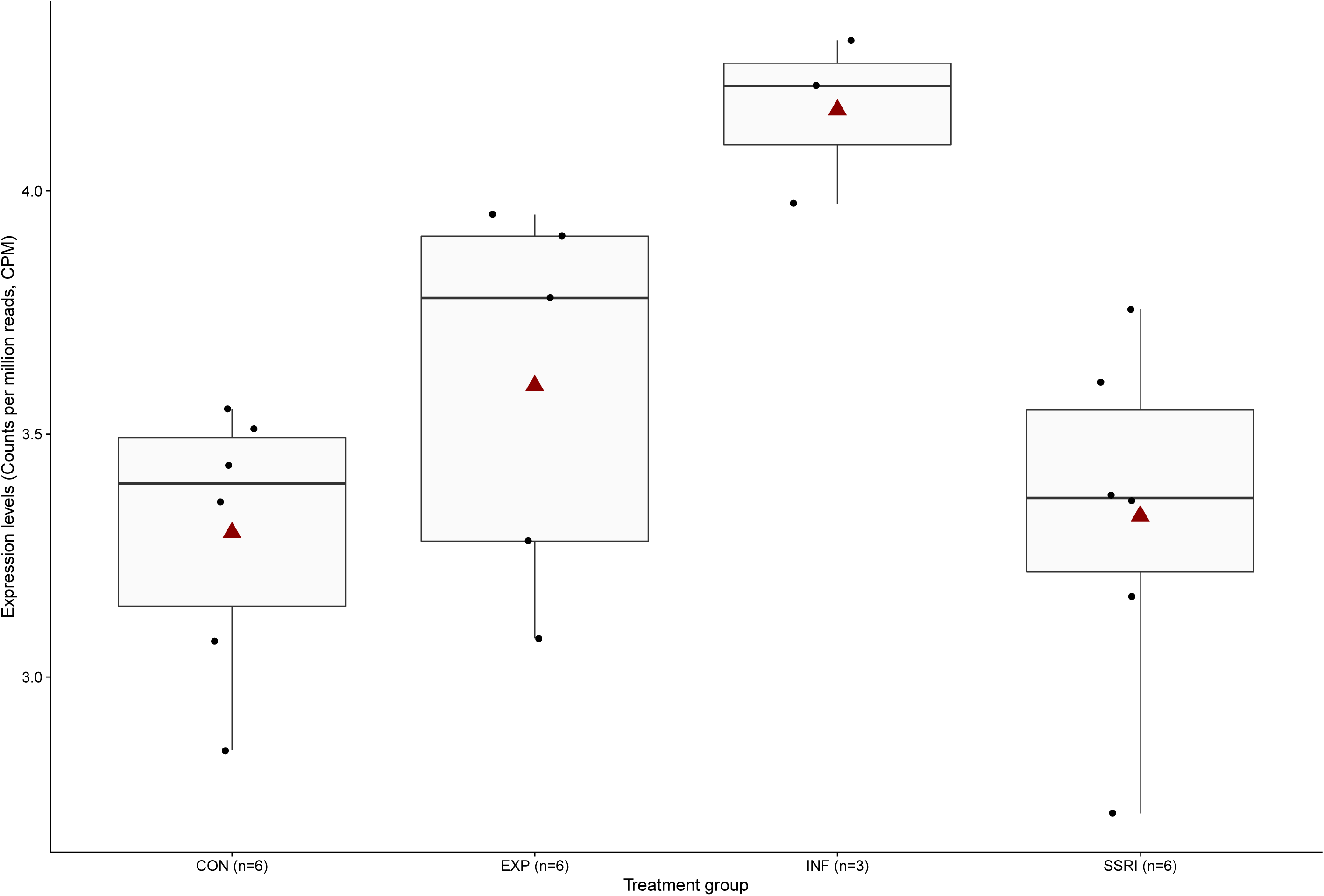
The IMPA 1 gene is significantly more expressed in the brain of *S*.*solidus*-infected sticklebacks compared to exposed, SSRI-treated and control fish. Box plots with median, 25th and 75th percentiles, vertical bars representing the largest value no further than 1.5 times the interquartile range, means (red triangles), and individual data (black dots). The four treatment groups are: CON = healthy controls, EXP = fish that were exposed to the parasite but did not become infected, INF = infected fish, SSRI = SSRI-treated fish, sample size in parentheses.

### Functional analysis in infected sticklebacks: pharmacological rescue of behaviour

Several genes coding for molecules involved in the inositol pathway were found to be differentially regulated in the brain of *S. solidus*-infected fish (see Results section). One of them is inositol monophosphatase 1 (IMPA1, table S1). This gene codes for the IMPAse 1 enzyme, which is a central step in the synthesis of myo-inositol [44]. Altered inositol metabolism has been implicated in various human neuropsychiatric and neurological diseases [45]. Lithium chloride is used to diminish behavioural symptoms of these diseases, such as the manic phase symptoms observed in bipolar patients [46], which include sleeplessness, hallucinations, psychosis, or paranoid rage [47]. One of the most accepted mechanisms of lithium action is the *inositol depletion hypothesis* [44] which suggest that lithium acts by blocking the IMPase 1 enzyme activity, leading to a depletion of inositol in the brain of treated patients [44, 45, 48, 49]. Therefore, we predicted that we could rescue normal behaviour in infected stickleback by modulating the inositol pathway using lithium exposure. We measured two well-characterized behaviours in infected individuals: the tendency to swim near the surface and the response to a simulated bird strike (here the time spent frozen after an attack). Using wild-caught threespine sticklebacks from Lac Témiscouata (QC, Canada), we quantified these two behaviours in *S*.*solidus*-infected individuals before and after exposure to lithium. One group was exposed to two low doses of lithium (at 2.5 mM and 5mM), and a second group to a high dose (at 15 mM). Significant effects on behaviour in treated infected fish were tested for each dose using a linear mixed effects analysis of the relationship between our dependent variable (behaviour) and treatment. See supplementary material for details.

## Results and discussion

### Altered molecular pathways in the brain following an infection by S. solidus

There were 105 differentially expressed genes between infected and control sticklebacks: 92 up-regulated and 13 down-regulated in infected, with a median log2 fold change of 0.60 (range: 0.26 to 2.76) and - 0.40 (range: −0.82 to −0.28) respectively (table S1). A total of 45 out of the 105 differentially expressed genes were differentially expressed only in the infected-control comparison and are thus considered as a genomic signature of infection (37 up-regulated, 8 down-regulated), table S1). The 45 genes forming the infected genomic signature were significantly enriched for categories associated with behaviour alterations (aromatic amino acid transport, thyroid hormone transport), host response to infection (catalase activity, oligosaccharyl transferase activity), and cellular growth (thymidylate kinase activity, dTDP biosynthetic process) (table S2).

#### i) Molecular changes associated with behavioural alteration

Aromatic amino acids include all the precursors to biogenic amines (dopamine, serotonin, and epinephrine), melatonin, and thyroid hormone. Several biogenic amines are related to behaviour variation [50] and this result is in accordance with the altered serotonin metabolism found in the brain of infected fish [22]. The thyroid hormone transport function was also overrepresented in genes differentially expressed in infected fish. Administration of thyroid hormone (thyroxine) in the *Schistosoma mansoni* host increased worm numbers and lead to the development of giant worms [51]. Thyroid hormone can also affect behaviour : treatments with thyroid hormones cause salmons to move to open water in daytime [52] and to change from a territorial phase to schooling phase during smelting [53]. Thus, increase of thyroid hormone transport might be beneficial for *S. solidus* growth and the completion of its cycle.

IMPA 1 (inositol monophosphatase 1) was among the up-regulated genes in the brain of infected fish that is associated with behaviour (figure 1, table S1). This gene encodes IMPase 1, a central enzyme in the inositol pathway [48], which is implicated in a diverse range of responses in the central nervous system [44, 54]. Alterations to this signalling pathway could be the cause of behaviour changes in infected sticklebacks. We tested the functional link between an increase in IMPA1 expression and behaviour by pharmacologically blocking IMPase 1 activity with lithium, which is used to treat symptoms of bipolar disorder by targeting IMPase activity [45, 49]. We attempted to rescue two behaviours that are altered in *S*.*solidus*-infected individuals: the tendency to swim closer to the surface and the lack of response to predator attacks. Infected sticklebacks exposed to low doses of lithium chloride (2.5 and 5 mM) did not reduce the proportion of time they spent swimming in the upper part of the aquarium compared to the control week (2.5 mM, t-ratio = 0.123, p = 0.99, n = 17; 5 mM, t-ratio = −0.607, p = 0.82, n = 17, figure 2a). However, infected sticklebacks treated with lithium chloride at a dose of 15 mM spent significantly less time in the top of the aquarium than before treatment (figure 2a) (t-ratio = 5.69, p < 0.001, n = 5). Infected sticklebacks treated with both low doses of lithium chloride spent almost no time frozen after a simulated bird strike, which was not significantly different from their behaviour before treatment (figure 2b) (2.5 mM of lithium, t-ratio = 0.742, p = 0.74, n = 17; 5 mM of lithium, t-ratio = −0.021, p = 0.99, n = 17). However, infected fish treated with lithium chloride at a dose of 15 mM spent significantly more time frozen after a simulated bird strike (figure 2b) (t-ratio = −2.803, p = 0.003, n = 5). These results suggest that lithium can block the IMPase 1 enzyme activity in the infected stickleback brain and alter their behaviour, making them respond more like healthy fish. Indeed, fish treated with high doses of lithium spent on average 7 % (sd = 7 %) of time near the surface and 85 seconds (sd = 62 seconds) frozen after a simulated bird strike, while non-infected sticklebacks studied in the same conditions in a separate study spent 6 % (sd = 6 %) of time near the surface and stayed frozen 34 sec (sd = 65 sec) (Alves and Aubin-Horth, unpublished). Such observations imply that alterations in the inositol pathway could be the direct cause, at least in part, of the striking behavioural alterations observed in this host-parasite model. To our knowledge, our results are one of the first examples of a pharmacological rescue of the behaviour of a host infected with a putative manipulative parasite, along with findings in the *Toxoplasma*-rodent system. Indeed, the risky behaviour of *Toxoplasma*-infected rodents towards predators has been successfully returned to cautiousness using antipsychotic drugs used to treat symptoms of schizophrenia [55].

**Figure 2.**
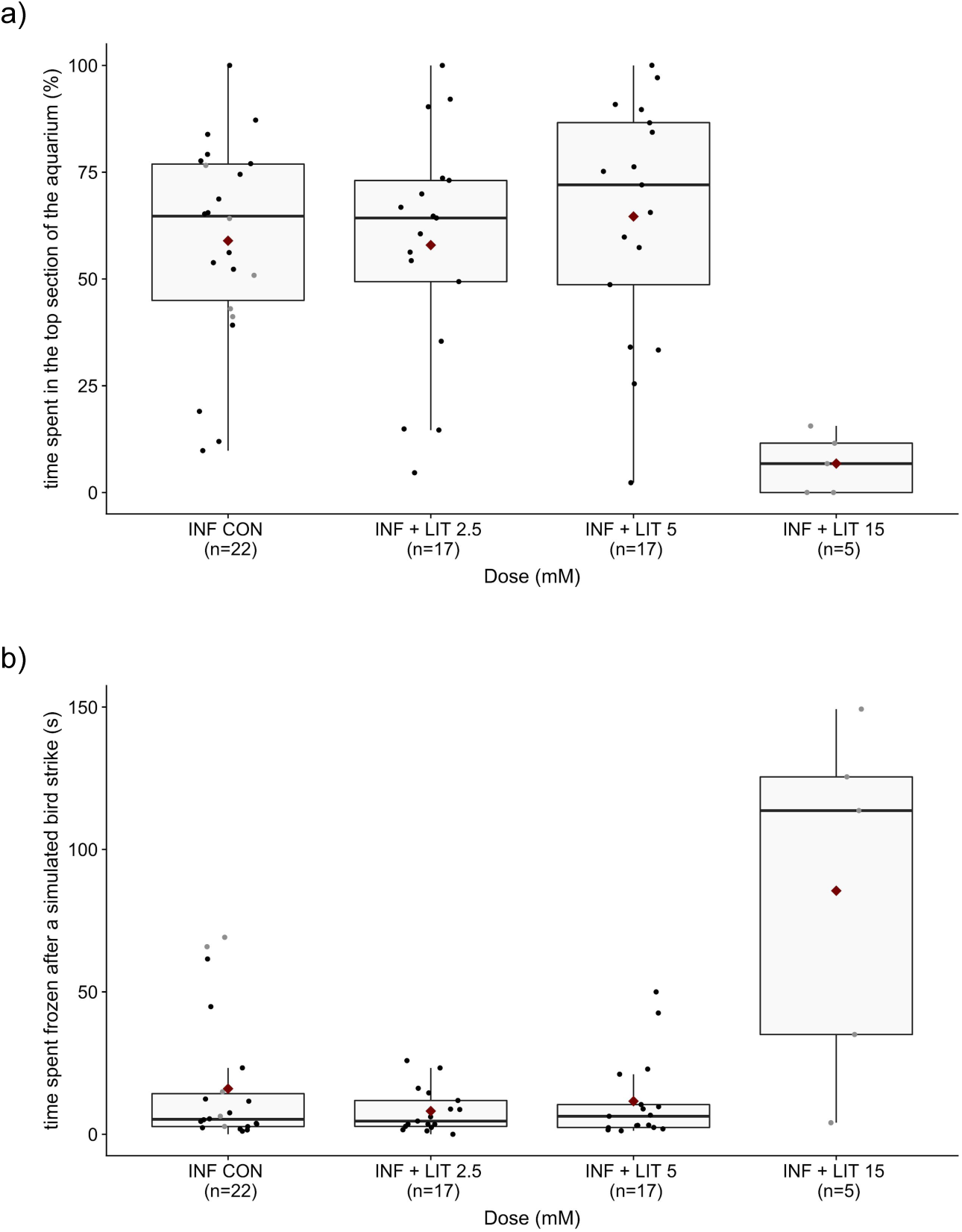
Lithium chloride exposure at a dose of 15mM rescues normal behaviours in *S*.*solidus*-infected sticklebacks. A) Infected sticklebacks treated with lithium chloride at a dose of 15 mM spent less time in the top of the aquarium compared to the control week (15 mM, t-ratio = 5.69, p = 0.0007, n = 5), while infected sticklebacks treated with lower doses of lithium chloride did not change their vertical preference after treatment (2.5 mM, t-ratio = 0.123, p = 0.99, n = 17; 5 mM, t-ratio = −0.607, p = 0.82, n = 17). B) Infected sticklebacks treated with lithium chloride at a dose of 15 mM spent more time frozen after a simulated bird strike in comparison to their control week (15 mM, t-ratio = −2.803, p = 0.003, n = 5), while infected sticklebacks treated with lower doses of lithium chloride did not change the time they spent frozen following a simulated bird strike after treatment (2.5 mM, t-ratio = 0.742, p = 0.74, n = 17; 5 mM, t-ratio = −0.021, p = 0.99, n = 17). Box plots with median, 25th and 75th percentiles, vertical bars representing the largest value no further than 1.5 times the interquartile range, means (red triangles), and individual data (black dots). The four treatment groups are: INF CON = infected control, INF LIT = infected fish treated with lithium chloride, with the number representing the dose in mM, sample size in parentheses.

In the *Schistocephalus*-stickleback system, it remains to be tested whether the alteration of the inositol pathway is a side-effect of a host response or if it is the result of a direct manipulation by the parasite. To test these different hypotheses, a combination of approaches will be needed to gather indirect and direct evidence [19]. Such approaches include characterizing which molecules are secreted/excreted by the parasite (termed “manipulation factors” [19], if these manipulation factors alter the host behaviour [56], and through which molecular mechanisms in the host.

#### ii) Molecular changes associated with the host response

Enrichment for certain biological functions in infected fish brains suggested that a host response to infection could be at play. The over representation of catalase activity, an important enzyme protecting the cell from oxidative damage by reactive oxygen species (ROS) [57] might be explained as a consequence of infection, since ROS production is increased in head kidney leucocytes in contact with *S. solidus* extracts *in vitro* [26] and appears to play an important role in stickleback defence against *S. solidus* [32]. However, the over-representation in catalase activity might also indicate a way by which the parasite can manipulate its host in order to eliminate an oxidative stress that would otherwise compromise parasite survival [58]. This over-representation could also indicate a reaction from the host aimed at decreasing the oxidative stress caused by the parasite. Genes whose function was associated with oligosaccharyl transferase activity were also over-represented in infected fish. This transferase is implicated in post-translational modifications of proteins by glycosylation, which determines the localization and function of these proteins [59]. Again, those modifications might be a global host response to infection at the protein level that can have major effects on cellular activity, among other wide-ranging effects. Finally, thymidylate kinase activity and dTDP biosynthetic process are biological functions enriched in infected fish brains that are related to cellular growth. The over representation of the thymidylate kinase activity, an important enzyme that assists biosynthesis of mitochondrial DNA, might also be explained as a consequence of infection, since a thymidylate kinase-like gene was found up-regulated in infected salmon and may be linked to the innate response to infection by a monogenean parasite [60]. An interesting feature of the *S. solidus*-stickleback system is that the hypothesis of a global host response could be verified by a gene expression study in the first intermediate host of *S. solidus*, the copepod. Determining if the biological functions of genes differentially expressed in infected copepod mirror the ones found in infected fish would allow us to determine the degree of overlap in the molecular response of both hosts and at the same time learn about the specificity of interactions at each life stage of the parasite [20, 61].

Our results come with limitations. Because of a small sample size, high biological variation within a group and small fold changes associated with the use of whole brain sampling, using a false discovery rate (FDR) resulted in little or no significant differentially expressed genes depending on the comparison. Genes found to be up- or down-regulated in INF fish will therefore each necessitate further functional validations, as for all gene expression studies that show an association between a phenotype and expression changes (rather than a causal link). Our test of a causal link between a disruption of the inositol pathway and behaviours typical of infected sticklebacks using a pharmacological treatment supports the notion that some of the genes differentially expressed in the brain of infected individuals are indeed associated with behaviour modification following infection. However, it is crucial to underscore that it is highly probable that most changes in gene expression do not affect behaviour. Some of them may control other changes observed in infected individuals at the physiological level, others may be related to a general host response to infection, while some of the changes in gene expression may be a side-product of the presence of the parasite, or simply be false positive. Once we confirm the causal role of a molecular pathway in behaviour variation, the next step would be to test which one, if any, of these pathways are directly manipulated by the parasite. In this host-parasite system, a next step would be to determine if the increase in IMPA1 activity is a direct manipulation by the parasite, an indirect effect of its presence, or an active response from the host [19].

### Overlap of exposed and infected fish transcriptomes

#### i) Overlap between exposed and infected fish

Contrary to our prediction, infected and exposed fish did not have similar brain expression profiles. Only nine genes were differentially expressed both in exposed and infected fish compared to controls (table S3). Interestingly, fold changes for these nine genes were very similar in amplitude between infected and exposed fish and were all in the same direction (see supp. Fig 2). Four of those nine genes were found as differentially expressed only in the comparison of each of these two treatments compared to control individuals: a solute carrier family protein, a myosin light chain, a lipase gastric, and a spermine acetyltransferase. On the other hand, only one gene was significantly differentially expressed between infected fish and exposed fish: gdpd5a, a glycerophosphodiester phosphodiesterase domain containing 5a, which was down-regulated in infected fish (FC=-0.418, p=0.002). This gene codes for a protein proposed (by similarity) to be involved in neurite formation, the regulation of the metabolite glycerophosphocholine (which can act as an osmolyte, https://hmdb.ca/metabolites/HMDB0000086, [62]), and in the cleavage of the GPI anchor of RECK, which in turns is involved in Wnt7-specific actions in the brain (https://www.uniprot.org/uniprot/Q8WTR4, [63]). Interestingly, a genomic study on *S. solidus* [64] has shown the existence of mimicry proteins (similar to the vertebrate host protein), with one of them belonging to the Wnt protein family and being the same protein found to be over-expressed in the head of orthopterans infected by a behaviour-altering hairworm [65]. Based on these similarities between two manipulating parasites and the observed change in the host, it could be proposed that a general disruption of cell-to-cell communication leading to various changes in behaviour may be at play [66].

#### ii) Exposed fish also have a distinct gene expression profile

Interestingly, the brains of exposed fish are also very different from the ones of healthy control fish. 153 genes were differentially expressed in exposed fish compared to control individuals (table S4): 48 up- and 105 down-regulated, with a median fold change of 0.34 (range: 0.22 to 1.43) and −0.47 (range: −1.16 to - 0.22), respectively. More than 85% of these genes were specific to that exposed vs control comparison, thus forming an “exposed” genomic signature, including 95 of the 105 down-regulated genes. The enrichment analysis (table S2) performed on the exposed-specific genes showed no significant enrichment. Whether this unique expression profile of exposed fish is a cause of the failure of the parasite to successfully infect these fish and was already present before infection, or if it is a long-lasting consequence of exposure (or both) is unknown. Indeed, we cannot exclude the fact that individuals that became infected and the ones that did not were already different before the experimental infection, as interaction between the genotype of the host and the parasite has been previously shown [67]. However, this is unlikely in our case, as all individuals come from the same crosses and laboratory environment. One way to test that these differences in expression existed before exposure and are the cause of the resistance to the parasite would be to redo a brain transcriptome analysis on a much higher number of control sticklebacks to detect difference in expression in the genes assigned to the exposed genomic signature in the present study. If so, it would suggest that the genotype of a proportion of individuals is the cause of differential gene expression rather than exposure to the parasite. On the other hand, being exposed is known to modify the immune system of the host. Sticklebacks resistant to parasitic eye flukes have a higher basic immunocompetence than more susceptible host [68]. A transcriptomic study of changes in gene expression in the head kidney following exposure to *S*.*solidus* in sticklebacks found that ROS production and recycling, B cell activation and targeting, and fibrosis appear to play important roles in defence against cestodes [32], but did not find significant differences in gene expression between exposed and control fish. Similarly, exposed but non-infected gilthead sea bream (*Sparus aurata*) appear closer to the unexposed fish than the infected fish in their gene expression [69].

It is worth noting that while successful infection resulted in the up-regulation of genes in the brain, exposure without infection resulted mostly in lower expression of genes in the brain, suggesting that they reflect different processes. The transcriptome response to pathogen lines of different virulence also show this opposite response in a *Daphnia* host, with the infective line resulting in more down-regulated genes, while the non-infective strain resulted in up-regulation of genes, with little overlap between the genes affected [70]. It thus appears that exposure could have a significant and distinct effect in the brain of the host, even when infection ultimately fails, and that this effect is carried over several weeks.

### Overlap of SSRI-treated and infected fish transcriptomes

The analysis of the overlap between SSRI and infected fish brain expression profiles revealed only four genes in common (table S5). These four genes are related to cellular organization and transport or have unknown functions. In comparison, changes in gene expression specific to the SSRI treatment (table S6) include genes that have functions related to neurotransmission. Drugs in the SSRI family are designed to target the serotonin axis, which has its own range of effects on behaviour and physiology. The small overlap between genes affected in infected individuals and in SSRI-treated ones may thus be explained in part by the fact that SSRIs are specifically designed for a unique target, while *S. solidus* affects several traits in addition to behaviour. Infections have other major impacts on host sticklebacks that include reduced body condition, changes in metabolism, nutrient balance, and reproduction [7]. A small overlap does not suggest necessarily that *S. solidus* and SSRI target different molecular pathways, only that these similarities are masked by multiple molecular differences.

## Conclusion

Parasitic alteration of animal behaviour is predicted to be caused by, and to result in, multiple physiological changes in the host. Our study aimed at determining the brain gene expression profiles of parasitised fish to identify a genomic signature that distinguishes a *S*.*solidus*-infected stickleback from a control fish. Obtaining a general portrait of molecular pathways affected in an infected host is a valuable step to help determine if these hosts are manipulated by their parasite. Indeed, once it is known what changes in the brain of a host, one can manipulate these molecular pathways to recreate the infected host phenotype (or parts of it) in healthy individuals and ultimately test evolutionary predictions about effects on the parasite’s fitness of host’s behavioural differences, as done in *Gammarus* [71]. Our results allow us to predict that if *S*.*solidus* directly manipulates its fish host, its manipulations factors could target the inositol pathway. A next step would thus be to determine if *S. solidus* directly alters the behaviour of its host by affecting the inositol pathway, and if it does, to test if sticklebacks with an experimentally-altered inositol pathway consequently exhibit altered behaviours that increase the probability of being predated by the final avian host of *Schistocephalus solidus*.

## Supporting information

RNA-seq dataset

Behaviour dataset

## Acknowledgments

We thank the IBIS genomics analysis platform for help with RNA extraction and library preparation, L. Benestan for the specific subsetting idea, and J. Turgeon, C. R. Landry and J. Le Luyer for discussion. Funding for this project was provided by the Fonds de Recherche du Québec Nature et Technologie (FRQ-NT) through a Team project grant to NAH, the Natural Sciences and Engineering Research Council of Canada Discovery grant to NAH, and a NSERC Vanier fellowship to FOH.

## Authors’ contributions

L.G. designed the study with the input of N.A.-H. and I.B. L.G. extracted RNA from the brains. L.G. and F.-O.H. performed transcriptomic and statistical analyses. VAA designed and performed the lithium pharmacological manipulation experiments. L.G. and N.A.-H. drafted the manuscript with input from F.-O.H., VAA, and I.B. All authors reviewed and gave input on the final version of the manuscript.

## Ethics

The experimental exposure to *S. solidus* parasites and / or SSRIs was undertaken at the University of Leicester, UK, under the authority of a UK Home Office project license (PPL 70/8148, held by I.B.). The lithium exposure was performed at Université Laval under the CPAUL certificate number 2017085-2. The project was authorised by the Comité de Protection des Animaux de l’Université Laval (2014069-1).

## Data availability

The RNA-seq dataset consists of raw read counts for the 17417 transcripts that passed quality control and filters. Behaviour data from the lithium treatment is presented for all individuals.

## Electronic supplementary material

A) Supplementary tables. Results of statistical analyses of differential gene expression, overlap of differentially expressed genes between two treatments, and results of enrichment analyses.

B) Supplementary figures.

C) Supplementary material and methods.

