## Supplemental methods for "Host behavior alteration by its parasite: from brain gene expression to functional test"

**Supporting Materials and Methods**

**Exposure of fish host to its parasite or SSRI**

All the threespine sticklebacks analysed in this study were the same samples used and described in [1]. We caught wild adults (mid-Wales, UK) and brought them into breeding condition. We reared the resulting fry for six months. Fish that were selected as hosts for the parasite were exposed to infected copepods. Each fish was starved for 48h to increase the probability that the fish successfully ingested the infected copepod. We isolated each fish in 1L (150 mm × 80 mm × 80 mm) plastic tanks, where they were each fed one infected copepod and then left for one week in their individual, filtered tank. We waited for three months for parasite growth. This treatment resulted in infected (fish with a parasite) and exposed fish (fish without a parasite). It was not possible to distinguish the exposed and infected individuals prior to dissection. The immersion protocol to obtain SSRI-treated fish was done in parallel (see [1]). Fluoxetine exposure decreases fish swimming velocity in medaka larvae and increases the time spent at the edges of aquaria at a concentration of 1mg/L for 72h [2]. We therefore used a fluoxetine concentration in the water of 1 mg/L (Fluoxetine HCl, BML-NS140, Enzo Life Sciences Inc., USA), which resulted in behavioural changes [1]. Control fish that were never exposed to a parasite were kept in the same conditions in parallel.

**Tissue sampling**

Fish were euthanized with a benzocaine (10 g/L) immersion. We measured the length and weight of the fish. One fish (SSRI treatment) died during the last 24 hours of treatment. Brains were rapidly dissected with RNase-zap-treated surgical pliers and placed individually in Eppendorf tubes containing 1mL of RNAlater (Ambion Inc., Austin, TX, USA), left at 4°C for 24h and then kept at -80 °C until later use.

**RNA preparation, library construction, and sequencing**

We extracted total RNA from the brain of six control, three infected, six exposed (but not infected) and six SSRI-treated fish (total of 21 individuals, all females) using a standard Trizol reagent protocol (miRNeasy Micro kit, Qiagen) and stored it at -80°C. Concentrations and 260/280 ratios were quantified by spectrophotometer (Nanodrop, Thermo scientifics) while RNA purity and quality were measured using the RNA 6000 Nano Kit (Agilent Technologies Inc., USA). We produced libraries for these 21 individuals using the TruSeq RNA Library Prep Kit v2 (Illumina, Inc., USA) with a unique barcode for each library. Library quality and size was assessed on a Bioanalyzer High Sensitivity DNA Assay (Agilent). The 21 cDNA libraries were then pooled before being sequenced on an Illumina HiSeq 2000 at the Génome Quebec Innovation Center (McGill University, Montréal, QC, Canada) using single-read technology (1x100 bp), for a total of three flow cell sequencing lanes. In total, 43 Gb of raw data was generated, which represents 594 million x 100 bp single-read sequences distributed across the 21 samples with an average size of 13.5 Mb (median = 13.5 Mb, range = 6.1-18.8 Mb).

**RNA-seq data analysis**

Quality control was first performed on all sequencing libraries using FastQC (Andrews 2010, http://.bioinformatics.babraham.ac.uk/projects/fastqc/). We then removed any remaining sequencing adaptors from raw reads using Trimmomatic v.0.33 [3], The libraries were then quality trimmed with the following settings (Phred score > 2, ILLUMINACLIP:2:30:10, SLIDINGWINDOW:20:2, LEADING:2, TRAILING:2) and size-selected (MINLEN:60).

We aligned all trimmed reads on the reference *Gasterosteus aculeatus* transcriptome (http://www.ensembl.org, version 83), allowing the estimation of transcript-specific expression levels for each individual fish. We aligned short reads from the 21 individual HiSeq libraries on the reference transcriptome using the BWA-MEM algorithm v.0.7.13 [4] with default parameters. The reference transcriptome contains in total 27,628 transcript sequences corresponding to actual and possible genes identified through the standard Ensembl genebuild pipeline. Reads with a unique hit on the reference were kept, while ambiguous multi-mapping reads were discarded from the final alignment. We obtained raw read counts for each transcript in each individual brain contained in SAM files using htseq-count v.0.6.1 [5] with default parameters. In total, we detected the expression of 26,073 transcripts (i.e. transcript-specific raw read count > 3 in at least 2 samples, representing 94% of the reference transcriptome). Transcript-specific read counts were used in downstream analyses as raw expression levels in a read count matrix. Official gene annotation for the *G. aculeatus* reference genome version 83 was downloaded from Ensembl and added as a column in the raw read-count matrix.

**Normalisation and filtering of transcripts**

We used the package ‘edgeR’ v.3.24.3 [6] available in the R software version 3.0.2 (R Foundation for Statistical Computing, Vienna, Austria) to filter and prepare raw read counts for differential expression analysis. As a first filter, we only kept transcripts that corresponded to annotated genes from *G. aculeatus* reference genome version 83. This resulted in 17,417 annotated transcripts. As a second filter, we kept transcript sequences that showed at least 15 Counts Per Million (CPM), in at least three samples for control , exposed and SSRI-treated groups, and in at least two samples for the infected fish group, using the cpm function of the edgeR package [6]. This filter retained in total 12,520 transcripts (45%). Read counts for these transcripts were then normalized with the “calcNormFactors” function using the Trimmed Mean of M values (TMM) method in order to correct for unequal library sizes [7].

**Library quality control**

We performed a Multidimensional Scaling Plot (MDS) with the function “plotMDS” from the R package ‘limma-voom’ v.3.38.1 [8] on the top 1000 most differentiated transcripts to detect potential outlier libraries and verify that samples from the same groups clustered together. This allowed the detection of an outlier library (exposed but not infected fish #20), which was then discarded from downstream analyses (labelled as EXP20 in supplementary figure 1). We performed statistical analyses on the remaining 20 individuals.

**Analysis of differential gene expression**

We used the R package “limma-voom” v.3.7 [8] to determine differential gene expression. Absolute read counts were converted into their respective CPM value and log2-transformed using the “voom” function from the limma package. Each transcript was fitted to an independent linear model using the log2(CPM) values as the response variable and the treatment (control, infected, SSRI-treated, exposed) as the explanatory variable. No intercept was used and comparisons of interests between treatments were performed. Each linear model was then analyzed through limma's Bayes pipeline. This last step allowed the discovery of differentially expressed transcripts based on a p-value of p<0.005 to determine which genes were differentially expressed.

**Defining genome-wide expression profiles of each treatment**

We quantified the number of genes that show changes in their expression profiles only in one specific treatment and which ones change in expression in two or more treatments. This allows us to define the genomic signature associated with each treatment, which we define as genes that are differentially regulated in one treatment compared to all other treatments, as well as overlaps between treatments to test some of our predictions.

**GO enrichment analysis**

The genomic signature also allows us to perform an enrichment analysis using the GO terms associated with genes that behave similarly within a treatment. We determined if certain biological functions were significantly over represented in a given group of genes with similar expression profiles using the Python package ‘goatools’ v.0.6.5 [9] to perform Fisher’s exact tests on GO annotation terms (parameters: --ident, --no_propagate_counts, with go-basic reference set v.1.2). We performed one enrichment test per group of genes composing each genomic signature. Specifically, we tested if GO terms were significantly over-represented among the subset of differentially expressed genes composing a given genomic signature (i.e. genes that are specifically differentially expressed in that particular treatment vs control comparison), as compared to all the expressed genes. Annotation of GO terms for each gene was extracted from the published transcriptome of *Gasterosteus aculeatus* and GO terms overrepresented in a given group, compared to the reference transcriptome, were analyzed. We used a p-value of p<0.005 as a threshold indicating over representation of a GO term.

**Functional analysis in infected stickleback**

*Fish sampling and rearing*

We caught threespine sticklebacks in September of 2017. All fish were part of a wild population from Lac Témiscouata, QC, Canada (47°40′33′′N, 68°50′15′′W, freshwater environment). We collected fish with a seine net and minnow traps. Before transportation, we kept fish in coolers with water from the lake. Air stones were constantly used to keep water aerated, including during transportation. We brought fish to the Laboratoire Aquatique de Recherche en Sciences Environnementales et Médicales (LARSEM) at Université Laval. Adults and juveniles were kept in separate tanks of 3 L, under natural light:dark photoperiod (13:11) and water temperature of 12 ºC. We fed fish every day, twice a day, with a mixture of brine shrimps (Hikari Bio-Pur) and flakes (Nutrafin-Basix). All work was carried out in compliance with Animal Care and Use Guidelines, under a permit of the Comité de Protection des Animaux de l’Université Laval (CPAUL, permit 2017-085-*2*).

*Selection of drug and pharmacological doses*

We used a pharmacological approach to test the hypothesis that the *myo*-inositol pathway is directly implicated in the behavioural alteration observed in infected sticklebacks. We aimed to reduce cerebral levels of endogenous *myo-*inositol of infected sticklebacks by blocking the IMPase 1 enzyme. To do so, we used lithium chloride, a drug used in bipolar disorder patients. Patients suffering from bipolar disorder have higher than normal activity of IMPase 1 [10], and higher cerebral *myo*-inositol levels during the manic phase [11]. The disease symptoms can be decreased by the intake of lithium, which inhibits IMPase 1 enzymatic activity and reduces inositol levels [10]. The most accepted hypotheses to explain lithium action is the *inositol depletion hypothesis*, based on the observation that lithium inhibits IMPase in vitro [12]. The inositol depletion hypothesis proposes that lithium interferes with the regeneration of inositol and, under conditions where inositol limits phosphatidylinositol (PI) synthesis, depletes the cell of PI [12]. No information on lithium doses in threespine sticklebacks was available, so we treated fish with lithium chloride by static exposures at concentrations of 2.5, 5, and 15 mM for three days, based on [13].

*Pharmacological manipulation experiment*

*1) 2.5 mM and 5 mM doses*

The first week was used as a control week to quantify baseline fish behaviour before treatment. Seventy-two hours before the behavioural tests, fish were transferred to a new individual aquarium of 2.7 L water provided with air stones (n = 17). After 48 hours, we changed all the water from each aquarium and we no longer fed fish until the day of the behavioural tests. Also, 24 hours before their behaviour was measured, we put each small aquarium inside a white plexiglass box to acclimate fish to white walls, since the behavioural tanks are made of white plexiglass. After 72h of acclimation, fish were submitted to behavioural tests.

After we performed all behavioural tests, we allowed fish to rest in their home tanks for three days where we fed them as usual (twice per day). After these three days, we transferred fish again to the new isolated tanks. Fish received a dose of 2.5 mM of lithium chloride (n = 17) each day, for three days. As before, we changed the water from each aquarium 48h after the transfer and no longer fed fish until the behavioural tests. We performed all steps described as above a third time (third week of tests) but we exposed fish to lithium chloride at a dose of 5 mM (n = 17). These fish were thus tested 3 times.

*2) 15 mM dose*

In a separate experiment, we performed the same procedure as described above for the control week and second week. We treated fish with lithium chloride at 15 mM (n = 5). These fish were thus tested 2 times.

*Behavioural tests*

We tested two behaviours: water depth preference and the response to a simulated bird predator attack. Infected individuals are found near the surface more often than non-infected sticklebacks [14] and infected individuals are bolder, recovering much faster from a predator attack than healthy ones [15].

*Water depth preference test*

For this test we used a tank (400 mm wide x 400 mm long x 600 mm height) made of three opaque walls (white Plexiglass) and one transparent front wall for video recording. The tank was filled with 500 mm of water. The water height was separated in three zones vertically. To stimulate fish activity, we placed a food bait (approx. 5 g of brine shrimps wrapped in a plastic film that was attached to a motor with a speed of 0.44 cm/s) in the upper third of the tank. An artificial plant was placed in the bottom of the aquarium as a refuge. Water was aerated continuously in order to keep the same oxygen concentrations in the upper and lower zones of the tank by an air stone. Before starting the test, fish were allowed to acclimate to this tank for five minutes. After this acclimation period, we tracked the fish vertical position in the tank for five minutes.

*Antipredator response*

For this test we used a different tank (300 mm wide x 600 mm long x 300 mm height) with all of its walls and floor internally coated with white Plexiglass. This tank was also equipped with a removable white wall that was used to create an acclimation zone of 180 x 295 mm. Before the beginning of each test we placed fish in this acclimation zone and after five minutes we gently removed the wall. We fed the fish with a plastic pipette, always putting the food item in front of its head. Once the fish started to eat, we performed a simulated bird strike from above using a model of a heron head (made of styrofoam and wood). If the individual did not eat, we waited a maximum of 5 minutes to perform the simulated bird strike. We analysed how long the fish spent frozen after the attack (not moving for at least more than 2 seconds).

All behavioural experiments were recorded using digital cameras (Super Circuits PC212XS) placed in front of the tank for the water depth preference test and on the top of the antipredator response test tank. Videos were later analysed using Ethovision software (Ethovision XT 11.5, Noldus Information Technology, Wageningen,[16]). Water depth preference tests were analysed with an automatic tracking module and time spent frozen was analysed using a manual behaviour setting module of the software.

*Statistical analysis*

We used R studio [17], R version 3.5.1 [18] and lme4 [19] to perform a linear mixed effects analysis of the relationship between our dependent variables (proportion of time spent in the upper zone of aquarium and time spent frozen after a simulated bird strike) and treatment. For the water depth preference test, we used the proportion of time spent in the upper part of the aquarium. Thereafter, we performed an arcsine square root transformation to this variable for the statistical analysis (data in Figure 2A is presented without the transformation).

For the first experiment, we used as fixed effects the treatment dose (0, 2.5 or 5 mM of lithium chloride), fish sex and the experimental block. We added fish identification (id) as a random effect since the same fish was tested either three times (for the low lithium doses) or twice (high lithium dose). For the second experiment, we used the treatment dose (0 or 15 mM of lithium chloride) and fish sex as fixed effects. Fish id was set as a random effect. Visual inspection of residual plots did not reveal any obvious deviations from homoscedasticity or normality for the tests.

**Ethics statement**

The experimental exposure to *S. solidus* parasites and / or SSRIs was undertaken at the University of Leicester, UK, under the authority of a UK Home Office project license (PPL 70/8148, held by I.B.). The lithium pharmacological manipulation was performed at Université Laval under the CPAUL certificate number 2017085-2. The project was authorised by the Comité de Protection des Animaux de l’Université Laval (CPAUL, experimental animal use permit, certificate number 2014069-1).
