## Supplemental tables for "Host behavior alteration by its parasite: from brain gene expression to functional test"

Figure S1. Multidimensional scaling analysis performed on the transcriptome of stickleback brains (Top 1000 abundant genes) from four different groups (control, CON; SSRI-treated, SSRI; exposed to *S. solidus* but uninfected, EXP; and infected to *S. solidus*, INF) showing that one individual (EXP20) was different from the rest of the individuals (removed from dataset afterwards due to its poor library quality).

-1

0

1

2

3

-0.5

0.0

0.5

Leading logFC dim 1

Leading logFC dim 2

CON04

CON05

CON06

CON07

CON09

CON10

EXP01

EXP02

EXP07

EXP12

EXP17

EXP20

INF03

INF01

INF02

SSRI01

SSRI02

SSRI04

SSRI05

SSRI06

SSRI12

Figure S2. Scatter plot representing the log fold changes of the nine genes differentially expressed both in infected and exposed fish compared to controls. Their fold changes had similar amplitude and direction (up-regulated or down-regulated in both treatments).

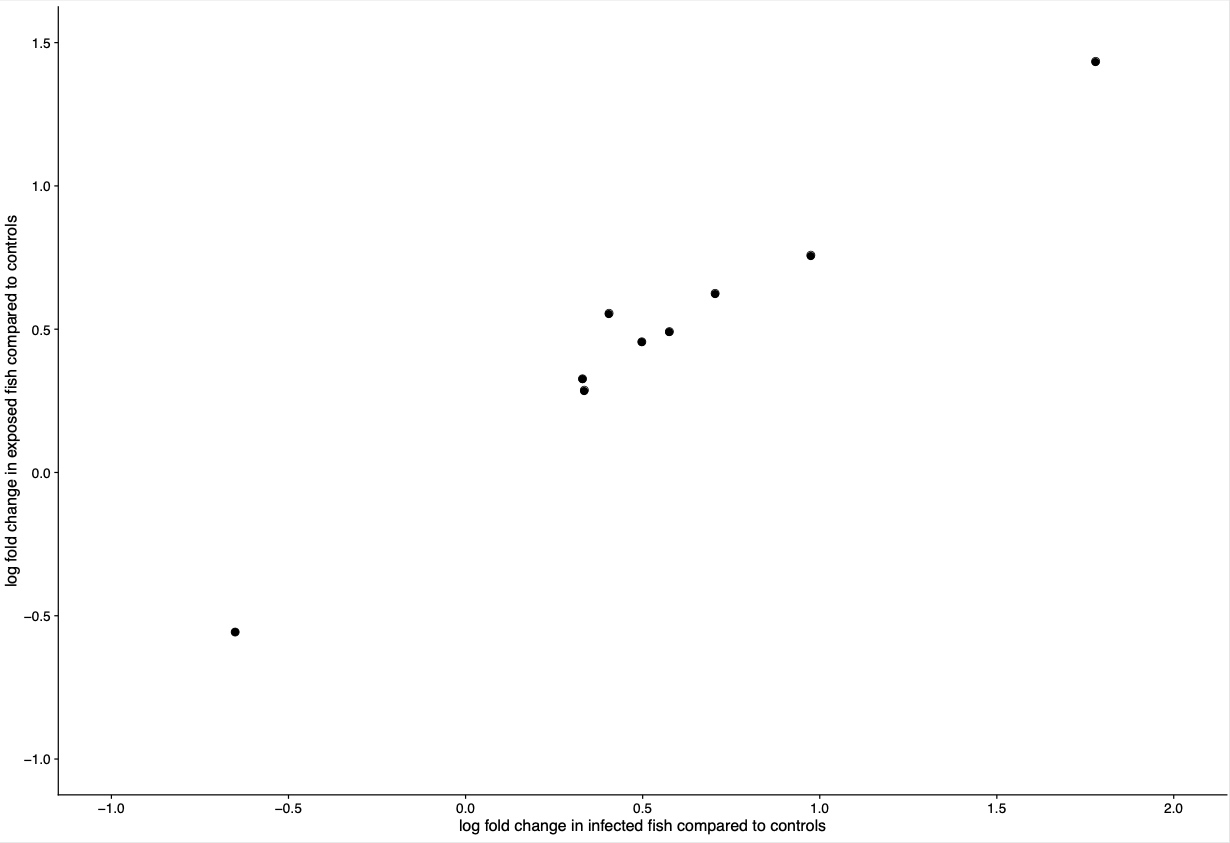
