## Supplemental figures for "Host behavior alteration by its parasite: from brain gene expression to functional test"

Table S1. Genes differentially expressed in the brain of *Schistocephalus* infected sticklebacks compared to control individuals. Infected individuals had 92 up-regulated and 13 down-regulated genes. Gene ID, Gene name and Annotation refer to the identity of the transcript according to the *Gasterosteus aculeatus* BROAD-S1 genome assembly, genebuild release #83, NCBI accession ID: GCA_000180675.1. Log-fold change in expression in infected fish compared to controls (logFC) and p-values (p-value) are presented. Genes forming the genomic signature of infected fish (genes differentially expressed only in this specific comparison) are in bold.

| **Gene ID** | **Gene Name** | **Annotation** | **logFC** | **p-value** |
| --- | --- | --- | --- | --- |
| **UP-REGULATED** | | | | |
| ENSGACT00000022963 | sytl4 | synaptotagmin-like 4 | 0.92 | 6.994E-05 |
| ENSGACT00000006510 | IMPA1 (3 of 3) | inositol(myo)-1(or 4)-monophosphatase 1 | 0.92 | 1.498E-04 |
| **ENSGACT00000005162** | **aldh9a1a.2** | **aldehyde dehydrogenase 9 family member A1a tandem duplicate 2** | **0.69** | **2.410E-04** |
| ENSGACT00000022970 | zdhhc9 | zinc finger DHHC-type containing 9 | 1.00 | 2.610E-04 |
| ENSGACT00000021906 | olfm2b | olfactomedin 2b | 0.72 | 3.103E-04 |
| ENSGACT00000020028 | mnx2b | motor neuron and pancreas homeobox 2b | 0.57 | 3.774E-04 |
| ENSGACT00000009046 | tom1 | target of myb1 membrane trafficking protein | 0.40 | 4.661E-04 |
| ENSGACT00000019796 | DPP10 (2 of 3) | dipeptidyl-peptidase 10 (non-functional) | 0.37 | 4.762E-04 |
| ENSGACT00000011412 | mast3b | microtubule associated serine/threonine kinase 3b | 0.33 | 5.019E-04 |
| ENSGACT00000015847 | rcbtb2 | regulator of chromosome condensation (RCC1) and BTB (POZ) domain containing protein 2 | 0.69 | 5.474E-04 |
| ENSGACT00000010644 | sfxn3 | sideroflexin 3 | 0.64 | 5.851E-04 |
| ENSGACT00000018807 | fmnl1a | formin-like 1a | 0.65 | 5.940E-04 |
| ENSGACT00000018950 | tnrc6a | trinucleotide repeat containing 6a | 1.82 | 6.064E-04 |
| **ENSGACT00000004798** | **pnpla7b** | **patatin-like phospholipase domain containing 7b** | **1.30** | **6.330E-04** |
| ENSGACT00000004969 | nqo1 (7 of 10) | NAD(P)H dehydrogenase quinone 1 | 0.86 | 7.697E-04 |
| ENSGACT00000017393 | las1l | LAS1-like ribosome biogenesis factor | 1.76 | 7.917E-04 |
| **ENSGACT00000000659** | **snx21** | **sorting nexin family member 21** | **0.42** | **8.472E-04** |
| **ENSGACT00000005142** | **atp6ap1la** | **ATPase H+ transporting lysosomal accessory protein 1-like a** | **0.63** | **9.078E-04** |
| ENSGACT00000025888 | txndc11 | thioredoxin domain containing 11 | 0.95 | 1.028E-03 |
| ENSGACT00000013219 | lipf | lipase gastric | 1.78 | 1.090E-03 |
| ENSGACT00000015391 | si:dkey-27n14.1 | si:dkey-27n14.1 | 0.70 | 1.148E-03 |
| ENSGACT00000022584 | clul1 | clusterin-like 1 (retinal) | 0.77 | 1.183E-03 |
| ENSGACT00000010829 | myl9b | myosin light chain 9b regulatory | 0.74 | 1.264E-03 |
| **ENSGACT00000004801** | **gata5** | **GATA binding protein 5** | **1.28** | **1.342E-03** |
| ENSGACT00000021804 | PITX2 (2 of 2) | paired-like homeodomain 2 | 2.76 | 1.349E-03 |
| ENSGACT00000011235 | txnl4b | thioredoxin-like 4B | 0.97 | 1.349E-03 |
| **ENSGACT00000025866** | **med21** | **mediator complex subunit 21** | **0.71** | **1.411E-03** |
| ENSGACT00000008004 | igf2bp2a | insulin-like growth factor 2 mRNA binding protein 2a | 0.67 | 1.479E-03 |
| ENSGACT00000026008 | sertad2a | SERTA domain containing 2a | 0.60 | 1.564E-03 |
| ENSGACT00000008144 | ghdc | GH3 domain containing | 0.41 | 1.600E-03 |
| ENSGACT00000025465 | mkrn1 | makorin ring finger protein 1 | 0.51 | 1.646E-03 |
| ENSGACT00000012825 | slc38a10 | solute carrier family 38 member 10 | 0.59 | 1.667E-03 |
| **ENSGACT00000004814** | **fstl1b** | **follistatin-like 1b** | **0.61** | **1.669E-03** |
| **ENSGACT00000006401** | **PRR12 (1 of 2)** | **proline rich 12** | **0.69** | **1.678E-03** |
| ENSGACT00000022942 | tnni2b.1 | troponin I type 2b (skeletal fast) tandem duplicate 1 | 0.54 | 1.683E-03 |
| ENSGACT00000004824 | klhl30 | kelch-like family member 30 | 0.64 | 1.738E-03 |
| ENSGACT00000015505 | ern1 | endoplasmic reticulum to nucleus signaling 1 | 0.63 | 1.779E-03 |
| ENSGACT00000026975 | CCHCR1 | coiled-coil alpha-helical rod protein 1 | 0.38 | 1.837E-03 |
| ENSGACT00000011070 | smtnb | smoothelin b | 0.50 | 1.895E-03 |
| ENSGACT00000021434 | apbb2b | amyloid beta (A4) precursor protein-binding family B member 2b | 0.29 | 1.942E-03 |
| **ENSGACT00000011059** | **rngtt (2 of 2)** | **RNA guanylyltransferase and 5'-phosphatase** | **0.63** | **2.125E-03** |
| **ENSGACT00000007465** | **rab22a** | **RAB22A member RAS oncogene family** | **0.37** | **2.159E-03** |
| **ENSGACT00000020085** | **GATAD2A (2 of 2)** | **GATA zinc finger domain containing 2A** | **1.23** | **2.225E-03** |
| ENSGACT00000025265 | afmid | arylformamidase | 1.49 | 2.235E-03 |
| **ENSGACT00000018172** | **mtrf1** | **mitochondrial translational release factor 1** | **0.33** | **2.499E-03** |
| **ENSGACT00000023496** | **NCSTN** | **nicastrin** | **0.93** | **2.551E-03** |
| **ENSGACT00000018927** | **ccl19a.1** | **chemokine (C-C motif) ligand 19a tandem duplicate 1** | **1.48** | **2.682E-03** |
| **ENSGACT00000001008** | **PRKCD (1 of 2)** | **protein kinase C delta** | **0.96** | **2.746E-03** |
| ENSGACT00000012323 | cep41 | centrosomal protein 41 | 0.71 | 2.751E-03 |
| ENSGACT00000022694 | slc35e4 | solute carrier family 35 member E4 | 0.47 | 2.778E-03 |
| ENSGACT00000011308 | hsd17b1 | hydroxysteroid (17-beta) dehydrogenase 1 | 0.71 | 2.795E-03 |
| **ENSGACT00000013462** | **fabp7a** | **fatty acid binding protein 7 brain a** | **0.53** | **2.931E-03** |
| ENSGACT00000026603 | amot | angiomotin | 0.62 | 2.948E-03 |
| ENSGACT00000005508 | DNM1L (1 of 2) | dynamin 1-like | 0.38 | 3.089E-03 |
| ENSGACT00000008453 | slc32a1 | solute carrier family 32 (GABA vesicular transporter) member 1 | 0.50 | 3.139E-03 |
| **ENSGACT00000004966** | **STT3B** | **STT3B subunit of the oligosaccharyltransferase complex (catalytic)** | **0.38** | **3.162E-03** |
| **ENSGACT00000009312** | **mapre3b** | **microtubule-associated protein RP/EB family member 3b** | **0.37** | **3.216E-03** |
| **ENSGACT00000002869** | **SLC16A3 (1 of 2)** | **solute carrier family 16 (monocarboxylate transporter) member 3** | **0.37** | **3.227E-03** |
| ENSGACT00000016723 | scap | SREBF chaperone | 0.81 | 3.243E-03 |
| **ENSGACT00000017293** | **zgc:152658** | **zgc:152658** | **0.37** | **3.362E-03** |
| **ENSGACT00000003359** | **zgc:109744** | **zgc:109744** | **0.74** | **3.459E-03** |
| ENSGACT00000013874 | oprl1 | opiate receptor-like 1 | 0.50 | 3.503E-03 |
| ENSGACT00000022029 | ankrd12 | ankyrin repeat domain 12 | 0.40 | 3.608E-03 |
| **ENSGACT00000005957** | **dtymk** | **deoxythymidylate kinase (thymidylate kinase)** | **0.42** | **3.665E-03** |
| **ENSGACT00000013209** | **cdh4** | **cadherin 4 type 1 R-cadherin (retinal)** | **1.59** | **3.681E-03** |
| **ENSGACT00000004407** | **sgca** | **sarcoglycan alpha** | **0.59** | **3.694E-03** |
| ENSGACT00000007951 | ncam3 | neural cell adhesion molecule 3 | 0.71 | 3.871E-03 |
| ENSGACT00000027061 | sat2b | spermidine/spermine N1-acetyltransferase family member 2b | 0.58 | 3.884E-03 |
| ENSGACT00000026691 | hnrpl | heterogeneous nuclear ribonucleoprotein L | 0.64 | 3.898E-03 |
| **ENSGACT00000002889** | **adarb1b** | **adenosine deaminase RNA-specific B1b** | **0.36** | **3.904E-03** |
| ENSGACT00000012274 | slc2a9l1 (1 of 2) | solute carrier family 2 (facilitated glucose transporter) member 9-like 1 | 0.41 | 4.019E-03 |
| **ENSGACT00000019698** | **ptbp2b (2 of 2)** | **polypyrimidine tract binding protein 2b** | **1.41** | **4.048E-03** |
| **ENSGACT00000010009** | **gstr (1 of 2)** | **glutathione S-transferase rho** | **0.26** | **4.048E-03** |
| ENSGACT00000007444 | osbpl2b | oxysterol binding protein-like 2b | 0.68 | 4.096E-03 |
| ENSGACT00000011406 | MYL2 (1 of 2) | myosin light chain 2 regulatory cardiac slow | 0.33 | 4.110E-03 |
| ENSGACT00000022412 | si:ch73-22a13.3 | si:ch73-22a13.3 | 0.40 | 4.229E-03 |
| **ENSGACT00000000121** | **rxrba** | **retinoid x receptor beta a** | **0.57** | **4.270E-03** |
| **ENSGACT00000018650** | **ints1** | **integrator complex subunit 1** | **2.37** | **4.408E-03** |
| **ENSGACT00000023753** | **GADD45G (3 of 3)** | **growth arrest and DNA-damage-inducible gamma** | **0.44** | **4.413E-03** |
| **ENSGACT00000004908** | **etv4** | **ets variant 4** | **0.33** | **4.549E-03** |
| **ENSGACT00000004808** | **g6pca.2** | **glucose-6-phosphatase a catalytic subunit tandem duplicate 2** | **0.39** | **4.568E-03** |
| ENSGACT00000012262 | tert | telomerase reverse transcriptase | 0.34 | 4.599E-03 |
| **ENSGACT00000006273** | **KIF9** | **kinesin family member 9** | **0.47** | **4.694E-03** |
| ENSGACT00000012822 | ctbs | chitobiase di-N-acetyl- | 0.80 | 4.713E-03 |
| ENSGACT00000026805 | ggnbp2 | gametogenetin binding protein 2 | 2.21 | 4.758E-03 |
| ENSGACT00000003187 | PYURF | PIGY upstream reading frame | 0.41 | 4.806E-03 |
| **ENSGACT00000003001** | **ZMIZ1 (1 of 2)** | **zinc finger MIZ-type containing 1** | **0.36** | **4.824E-03** |
| **ENSGACT00000018093** | **xrcc3** | **X-ray repair complementing defective repair in Chinese hamster cells 3** | **0.46** | **4.824E-03** |
| **ENSGACT00000016661** | **spag9a** | **sperm associated antigen 9a** | **0.33** | **4.826E-03** |
| **ENSGACT00000022654** | **slc35c1** | **solute carrier family 35 (GDP-fucose transporter) member C1** | **0.49** | **4.841E-03** |
| ENSGACT00000004964 | fam160b1 | family with sequence similarity 160 member B1 | 0.34 | 4.967E-03 |
| ENSGACT00000017670 | ppp1r27a | protein phosphatase 1 regulatory subunit 27a | 0.52 | 4.980E-03 |
| **DOWN-REGULATED** | | | | |
| ENSGACT00000010617 | tns2b | tensin 2b | -0.82 | 2.092E-04 |
| **ENSGACT00000016231** | **lrrc73** | **leucine rich repeat containing 73** | **-0.41** | **1.192E-03** |
| ENSGACT00000018666 | ckap5 | cytoskeleton associated protein 5 | -0.65 | 1.591E-03 |
| **ENSGACT00000008616** | **slc16a10** | **solute carrier family 16 (aromatic amino acid transporter) member 10** | **-0.40** | **2.043E-03** |
| **ENSGACT00000024563** | **POU3F4** | **POU class 3 homeobox 4** | **-0.43** | **2.333E-03** |
| ENSGACT00000004216 | slc7a5 | solute carrier family 7 (amino acid transporter light chain L system) member 5 | -0.30 | 2.525E-03 |
| **ENSGACT00000012809** | **si:dkey-205k8.5** | **si:dkey-205k8.5** | **-0.52** | **3.556E-03** |
| **ENSGACT00000019385** | **ifi35** | **interferon-induced protein 35** | **-0.42** | **3.765E-03** |
| ENSGACT00000020268 | cryaa | crystallin alpha A | -0.30 | 3.961E-03 |
| **ENSGACT00000004596** | **cat** | **catalase** | **-0.38** | **4.393E-03** |
| **ENSGACT00000022490** | **apoda.2 (2 of 3)** | **apolipoprotein Da duplicate 2** | **-0.38** | **4.499E-03** |
| **ENSGACT00000006733** | **smad6b** | **SMAD family member 6b** | **-0.31** | **4.512E-03** |
| ENSGACT00000004211 | naa50 | N(alpha)-acetyltransferase 50 NatE catalytic subunit | -0.28 | 4.513E-03 |

Table S2. Biological functions over-represented in genomic signatures of a specific comparison between two groups. We used an enrichment analysis using Gene Ontology to detect GO terms significantly enriched in a given signature with a cut-off of p < 0.005. GO terms and GO descriptions were extracted from the output of the Python package ‘goatools’. The number of genes in the signature related to a GO term compared to the total number of genes in the genomic signature (Ratio) and the p-value (p-value) are presented.

| **Comparison** | **GO term** | **GO description** | **Ratio** | **p-value** |
| --- | --- | --- | --- | --- |
| INFECTED VS CONTROLS | GO:0015801 | aromatic amino acid transport | 1/45 | 4.59E-03 |
|  | GO:0070327 | thyroid hormone transport | 1/45 | 4.59E-03 |
|  | GO:0004096 | catalase activity | 1/45 | 4.59E-03 |
|  | GO:0015173 | aromatic amino acid transmembrane transporter activity | 1/45 | 4.59E-03 |
|  | GO:0004798 | thymidylate kinase activity | 1/45 | 4.59E-03 |
|  | GO:0006233 | dTDP biosynthetic process | 1/45 | 4.59E-03 |
|  | GO:0015349 | thyroid hormone transmembrane transporter activity | 1/45 | 4.59E-03 |
|  | GO:0004576 | oligosaccharyl transferase activity | 1/45 | 4.59E-03 |
| SSRI-TREATED vs CONTROLS | GO:0016998 | cell wall macromolecule catabolic process | 1/33 | 3.30E-03 |

Table S3. Genes that were significantly differentially expressed in both infected and exposed fish compared to controls. Nine genes were found using p-values with a cut-off of 0.005 to identify significantly differentially expressed genes in each comparison. Gene ID, Gene name and Annotation refer to the identity of the transcript according to the *Gasterosteus aculeatus* BROAD-S1 genome assembly, genebuild release #83, NCBI accession ID: GCA_000180675.1. Log-fold change in expression in infected fish compared to controls or exposed fish compared to controls (logFC) and p-values (p-value) are presented. Genes differentially expressed compared to controls only in these two comparisons are in bold.

|  |  |  | **INFECTED vs CONTROLS** | | **EXPOSED vs CONTROLS** | |
| --- | --- | --- | --- | --- | --- | --- |
| **Gene ID** | **Gene Name** | **Annotation** | **logFC** | **p-value** | **logFC** | **p-value** |
| ENSGACT00000008453 | slc32a1 | solute carrier family 32 (GABA vesicular transporter) member 1 | 0.50 | 3.139E-03 | 0.46 | 2.155E-03 |
| ENSGACT00000011235 | txnl4b | thioredoxin-like 4B | 0.97 | 1.349E-03 | 0.76 | 4.253E-03 |
| **ENSGACT00000011406** | **MYL2 (1 of 2)** | **myosin light chain 2 regulatory cardiac slow** | **0.33** | **4.110E-03** | **0.33** | **1.461E-03** |
| ENSGACT00000012262 | tert | telomerase reverse transcriptase | 0.33 | 4.599E-03 | 0.29 | 4.819E-03 |
| **ENSGACT00000012274** | **slc2a9l1 (1 of 2)** | **solute carrier family 2 (facilitated glucose transporter) member 9-like 1** | **0.41** | **4.019E-03** | **0.56** | **4.600E-05** |
| **ENSGACT00000013219** | **lipf** | **lipase gastric** | **1.78** | **1.089E-03** | **1.43** | **4.247E-03** |
| ENSGACT00000015391 | si:dkey-27n14.1 | si:dkey-27n14.1 | 0.70 | 1.148E-03 | 0.62 | 1.116E-03 |
| ENSGACT00000018666 | ckap5 | cytoskeleton associated protein 5 | -0.65 | 1.590E-03 | -0.56 | 1.421E-03 |
| **ENSGACT00000027061** | **sat2b** | **spermidine/spermine N1-acetyltransferase family member 2b** | **0.58** | **3.883E-03** | **0.49** | **4.429E-03** |

Table S4. Genes differently expressed in the brain of *Schistocephalus* exposed but non infected sticklebacks compared to controls. Exposed individuals had 48 up-regulated and 105 down-regulated genes using p-values with a cut-off of 0.005 to identify significantly differentially expressed genes. Gene ID, Gene name and Annotation refer to the identity of the transcript according to the *Gasterosteus aculeatus* BROAD-S1 genome assembly, genebuild release #83, NCBI accession ID: GCA_000180675.1. Log-fold change in expression in exposed fish compared to controls (logFC) and p-values (p-value) are presented. Genes forming the genomic signature of exposed fish (genes differentially expressed only in this specific comparison) are in bold.

| **Gene ID** | **Gene Name** | **Annotation** | **logFC** | **p-value** |
| --- | --- | --- | --- | --- |
| **UP-REGULATED** | | | | |
| ENSGACT00000012274 | slc2a9l1 (1 of 2) | solute carrier family 2 (facilitated glucose transporter) member 9-like 1 | 0.56 | 4.601E-05 |
| ENSGACT00000021266 | polr1e | polymerase (RNA) I polypeptide E | 0.32 | 6.555E-04 |
| ENSGACT00000009718 | sox10 (1 of 2) | SRY (sex determining region Y)-box 10 | 0.35 | 6.697E-04 |
| ENSGACT00000015105 | itih6 | inter-alpha-trypsin inhibitor heavy chain family member 6 | 0.29 | 8.604E-04 |
| ENSGACT00000018860 | gsx2 | GS homeobox 2 | 0.46 | 8.829E-04 |
| ENSGACT00000008458 | nptna | neuroplastin a | 0.45 | 9.211E-04 |
| ENSGACT00000020971 | vrk3 | vaccinia related kinase 3 | 0.51 | 9.276E-04 |
| ENSGACT00000014669 | ptprjb.2 | protein tyrosine phosphatase receptor type Jb tandem duplicate 2 | 0.33 | 1.066E-03 |
| ENSGACT00000006929 | kcna2b | potassium voltage-gated channel shaker-related subfamily member 2b | 0.37 | 1.082E-03 |
| ENSGACT00000015391 | si:dkey-27n14.1 | si:dkey-27n14.1 | 0.63 | 1.116E-03 |
| ENSGACT00000020832 | si:dkey-163m14.2 | si:dkey-163m14.2 | 0.47 | 1.132E-03 |
| ENSGACT00000016213 | ecm1b | extracellular matrix protein 1b | 0.42 | 1.237E-03 |
| ENSGACT00000024598 | gpr174 | G protein-coupled receptor 174 | 0.36 | 1.458E-03 |
| ENSGACT00000011406 | MYL2 (1 of 2) | myosin light chain 2 regulatory cardiac slow | 0.33 | 1.461E-03 |
| ENSGACT00000017452 | ikbkg | inhibitor of kappa light polypeptide gene enhancer in B-cells kinase gamma | 0.26 | 1.704E-03 |
| ENSGACT00000018634 | chsy1 | chondroitin sulfate synthase 1 | 0.28 | 1.856E-03 |
| ENSGACT00000015291 | hdac12 | histone deacetylase 12 | 0.24 | 2.095E-03 |
| ENSGACT00000008453 | slc32a1 | solute carrier family 32 (GABA vesicular transporter) member 1 | 0.46 | 2.155E-03 |
| ENSGACT00000024988 | CBLN1 (2 of 2) | cerebellin 1 precursor | 0.41 | 2.813E-03 |
| ENSGACT00000016204 | FBXO40 (1 of 3) | F-box protein 40 | 0.50 | 2.869E-03 |
| ENSGACT00000014341 | pmp22a | peripheral myelin protein 22a | 0.31 | 2.932E-03 |
| ENSGACT00000019356 | TRIM35 (23 of 27) | tripartite motif containing 35 | 0.56 | 3.024E-03 |
| ENSGACT00000018496 | dzip1 | DAZ interacting zinc finger protein 1 | 0.25 | 3.028E-03 |
| **ENSGACT00000007003** | **sorl1** | **sortilin-related receptor L(DLR class) A repeats containing** | **0.42** | **3.043E-03** |
| ENSGACT00000019789 | TUBA4A (2 of 2) | tubulin alpha 4a | 0.43 | 3.056E-03 |
| ENSGACT00000008146 | actc1a | actin alpha cardiac muscle 1a | 0.25 | 3.097E-03 |
| ENSGACT00000011928 | ZMAT1 | zinc finger matrin-type 1 | 0.25 | 3.170E-03 |
| ENSGACT00000007714 | CASKIN1 (1 of 2) | CASK interacting protein 1 | 0.24 | 3.215E-03 |
| ENSGACT00000021567 | sh3pxd2aa | SH3 and PX domains 2Aa | 0.27 | 3.234E-03 |
| ENSGACT00000008647 | adcy2a | adenylate cyclase 2a | 0.30 | 3.396E-03 |
| ENSGACT00000021551 | nip7 | NIP7 nucleolar pre-rRNA processing protein | 0.56 | 3.425E-03 |
| ENSGACT00000003298 | ZBTB22 (2 of 2) | zinc finger and BTB domain containing 22 | 0.28 | 3.439E-03 |
| ENSGACT00000021493 | si:ch1073-82l19.1 | si:ch1073-82l19.1 | 0.38 | 3.453E-03 |
| ENSGACT00000011774 | nacad | NAC alpha domain containing | 0.33 | 3.555E-03 |
| ENSGACT00000010676 | si:ch211-256m1.8 (1 of 4) | si:ch211-256m1.8 | 0.27 | 3.562E-03 |
| ENSGACT00000008661 | RPL27A | ribosomal protein L27a | 0.34 | 3.573E-03 |
| ENSGACT00000013982 | ncl1 | nicalin | 0.37 | 3.685E-03 |
| ENSGACT00000008544 | tmem205 | transmembrane protein 205 | 0.40 | 4.070E-03 |
| ENSGACT00000008420 | fam133b | family with sequence similarity 133 member B | 0.31 | 4.127E-03 |
| ENSGACT00000013219 | lipf | lipase gastric | 1.43 | 4.248E-03 |
| ENSGACT00000011235 | txnl4b | thioredoxin-like 4B | 0.76 | 4.253E-03 |
| ENSGACT00000008462 | SLC44A2 (1 of 2) | solute carrier family 44 (choline transporter) member 2 | 0.31 | 4.299E-03 |
| ENSGACT00000012296 | cacnb1 | calcium channel voltage-dependent beta 1 subunit | 0.23 | 4.325E-03 |
| ENSGACT00000013412 | prox1a | prospero homeobox 1a | 0.33 | 4.356E-03 |
| ENSGACT00000027061 | sat2b | spermidine/spermine N1-acetyltransferase family member 2b | 0.49 | 4.430E-03 |
| ENSGACT00000011921 | irx4b | iroquois homeobox 4b | 0.27 | 4.459E-03 |
| ENSGACT00000012262 | tert | telomerase reverse transcriptase | 0.29 | 4.820E-03 |
| ENSGACT00000015802 | slc7a4 | solute carrier family 7 member 4 | 0.27 | 4.493E-03 |
| **DOWN-REGULATED** | | | | |
| ENSGACT00000025269 | ino80b | INO80 complex subunit B | -0.64 | 8.306E-06 |
| ENSGACT00000017176 | MATN3 (2 of 2) | matrilin 3 | -0.61 | 1.585E-04 |
| **ENSGACT00000003485** | **fkbp7** | **FK506 binding protein 7** | **-0.70** | **1.604E-04** |
| ENSGACT00000026400 | chrnb1 | cholinergic receptor nicotinic beta 1 (muscle) | -0.72 | 1.749E-04 |
| ENSGACT00000016400 | chmp7 | charged multivesicular body protein 7 | -0.52 | 2.117E-04 |
| ENSGACT00000006708 | rpain | RPA interacting protein | -0.97 | 2.861E-04 |
| **ENSGACT00000017187** | **ube2e2** | **ubiquitin-conjugating enzyme E2E 2** | **-0.45** | **2.899E-04** |
| **ENSGACT00000008560** | **arhgef16** | **Rho guanine nucleotide exchange factor (GEF) 16** | **-0.43** | **3.012E-04** |
| ENSGACT00000002934 | slc16a5a (1 of 2) | solute carrier family 16 (monocarboxylate transporter) member 5a | -0.49 | 3.396E-04 |
| ENSGACT00000026668 | osr1 | odd-skipped related transciption factor 1 | -0.76 | 4.183E-04 |
| ENSGACT00000000642 | ARL4D | ADP-ribosylation factor-like 4D | -0.74 | 4.189E-04 |
| ENSGACT00000015014 | hrasb | #NAME? | -0.43 | 4.520E-04 |
| ENSGACT00000025370 | phyh | phytanoyl-CoA 2-hydroxylase | -0.46 | 5.038E-04 |
| ENSGACT00000002929 | taar15 (10 of 26) | trace amine associated receptor 15 | -0.44 | 6.346E-04 |
| ENSGACT00000013148 | notch3 | notch 3 | -0.39 | 6.459E-04 |
| ENSGACT00000019656 | tubgcp3 | tubulin gamma complex associated protein 3 | -0.34 | 6.499E-04 |
| ENSGACT00000018546 | bves | blood vessel epicardial substance | -0.75 | 8.266E-04 |
| ENSGACT00000017271 | txnrd2 | thioredoxin reductase 2 | -0.57 | 8.725E-04 |
| ENSGACT00000024423 | atp6v0e1 | ATPase H+ transporting lysosomal V0 subunit e1 | -0.51 | 8.840E-04 |
| ENSGACT00000013776 | si:ch73-280o22.2 | si:ch73-280o22.2 | -0.49 | 8.883E-04 |
| ENSGACT00000015380 | pmelb | premelanosome protein b | -0.33 | 9.056E-04 |
| ENSGACT00000011606 | crygm6 | crystallin gamma M6 | -0.51 | 9.403E-04 |
| ENSGACT00000014004 | mblac2 | metallo-beta-lactamase domain containing 2 | -0.35 | 9.470E-04 |
| ENSGACT00000027617 | wbscr16 | Williams-Beuren syndrome chromosome region 16 homolog (human) | -0.57 | 9.697E-04 |
| ENSGACT00000016574 | zgc:162816 | zgc:162816 | -0.64 | 1.120E-03 |
| ENSGACT00000021088 | tcf25 | transcription factor 25 (basic helix-loop-helix) | -0.51 | 1.156E-03 |
| ENSGACT00000011906 | ARHGEF25 (2 of 2) | Rho guanine nucleotide exchange factor (GEF) 25 | -0.61 | 1.361E-03 |
| ENSGACT00000018666 | ckap5 | cytoskeleton associated protein 5 | -0.56 | 1.421E-03 |
| ENSGACT00000004767 | cyc1 | cytochrome c-1 | -0.49 | 1.447E-03 |
| ENSGACT00000006635 | foxf2a | forkhead box F2a | -0.49 | 1.545E-03 |
| **ENSGACT00000000322** | **ints6** | **integrator complex subunit 6** | **-1.16** | **1.629E-03** |
| ENSGACT00000022352 | mgat4b | mannosyl (alpha-13-)-glycoprotein beta-14-N-acetylglucosaminyltransferase isozyme B | -0.41 | 1.672E-03 |
| ENSGACT00000023945 | stxbp1a | syntaxin binding protein 1a | -0.59 | 1.747E-03 |
| ENSGACT00000018367 | nup93 | nucleoporin 93 | -0.57 | 1.810E-03 |
| ENSGACT00000027408 | zgc:175280 (2 of 2) | zgc:175280 | -0.22 | 1.834E-03 |
| ENSGACT00000014475 | dazap2 | DAZ associated protein 2 | -0.31 | 1.864E-03 |
| ENSGACT00000018916 | adgrg1 (1 of 2) | adhesion G protein-coupled receptor G1 | -0.54 | 1.984E-03 |
| ENSGACT00000026572 | meig1 | meiosis/spermiogenesis associated 1 | -0.41 | 2.012E-03 |
| ENSGACT00000000060 | TRIM14 (2 of 74) | tripartite motif containing 14 | -0.65 | 2.076E-03 |
| ENSGACT00000013456 | ryr2b | ryanodine receptor 2b (cardiac) | -0.26 | 2.108E-03 |
| **ENSGACT00000002675** | **cherp** | **calcium homeostasis endoplasmic reticulum protein** | **-0.70** | **2.124E-03** |
| ENSGACT00000013819 | phc1 | polyhomeotic homolog 1 | -0.84 | 2.230E-03 |
| ENSGACT00000002817 | si:ch73-74h11.1 | si:ch73-74h11.1 | -0.60 | 2.238E-03 |
| ENSGACT00000020140 | ptges3b | prostaglandin E synthase 3b (cytosolic) | -0.78 | 2.281E-03 |
| ENSGACT00000021527 | slc24a3 (2 of 2) | solute carrier family 24 (sodium/potassium/calcium exchanger) member 3 | -0.27 | 2.312E-03 |
| ENSGACT00000000594 | METTL21B | methyltransferase like 21B | -0.26 | 2.363E-03 |
| ENSGACT00000007862 | ano3 | anoctamin 3 | -0.84 | 2.375E-03 |
| **ENSGACT00000002356** | **si:ch211-114l13.9 (9 of 22)** | **si:ch211-114l13.9** | **-0.46** | **2.389E-03** |
| ENSGACT00000006416 | ap5s1 | adaptor-related protein complex 5 sigma 1 subunit | -0.49 | 2.406E-03 |
| ENSGACT00000020070 | gtf3c3 | general transcription factor IIIC polypeptide 3 | -0.44 | 2.450E-03 |
| ENSGACT00000000297 | mfng | MFNG O-fucosylpeptide 3-beta-N-acetylglucosaminyltransferase | -0.43 | 2.473E-03 |
| ENSGACT00000009089 | col2a1b | collagen type II alpha 1b | -0.29 | 2.495E-03 |
| ENSGACT00000004950 | nqo1 (3 of 10) | NAD(P)H dehydrogenase quinone 1 | -0.54 | 2.555E-03 |
| ENSGACT00000022358 | rxfp3.3a1 | relaxin/insulin-like family peptide receptor 3.3a1 | -0.46 | 2.570E-03 |
| ENSGACT00000025810 | LRRC8E | leucine rich repeat containing 8 family member E | -0.42 | 2.618E-03 |
| ENSGACT00000013695 | kcnh5a | potassium voltage-gated channel subfamily H (eag-related) member 5a | -0.38 | 2.635E-03 |
| ENSGACT00000026473 | srfa | serum response factor a | -0.46 | 2.647E-03 |
| **ENSGACT00000000352** | **plxna3** | **plexin A3** | **-0.44** | **2.761E-03** |
| ENSGACT00000024870 | col4a5 | collagen type IV alpha 5 (Alport syndrome) | -0.68 | 2.765E-03 |
| ENSGACT00000022789 | slc20a1b | solute carrier family 20 (phosphate transporter) member 1b | -0.54 | 2.783E-03 |
| ENSGACT00000013939 | LRRC74A (2 of 2) | leucine rich repeat containing 74A | -0.37 | 2.790E-03 |
| ENSGACT00000008814 | atrnl1b | attractin-like 1b | -0.49 | 2.826E-03 |
| ENSGACT00000027711 | KIF2A (3 of 3) | kinesin heavy chain member 2A | -0.64 | 2.834E-03 |
| ENSGACT00000023114 | lef1 | lymphoid enhancer-binding factor 1 | -0.29 | 2.886E-03 |
| ENSGACT00000002412 | cpne3 (3 of 4) | copine III | -0.61 | 3.019E-03 |
| ENSGACT00000022343 | DGKQ | diacylglycerol kinase theta 110kDa | -0.34 | 3.031E-03 |
| ENSGACT00000006217 | mettl22 | methyltransferase like 22 | -0.81 | 3.059E-03 |
| ENSGACT00000001745 | TRIM35 (5 of 27) | tripartite motif containing 35 | -0.45 | 3.070E-03 |
| ENSGACT00000003551 | rttn | rotatin | -0.59 | 3.073E-03 |
| ENSGACT00000000612 | cotl1 | coactosin-like F-actin binding protein 1 | -0.44 | 3.201E-03 |
| ENSGACT00000011441 | crybb3 | crystallin beta B3 | -0.32 | 3.315E-03 |
| **ENSGACT00000002536** | **nup98** | **nucleoporin 98** | **-0.53** | **3.320E-03** |
| ENSGACT00000001495 | necap2 | NECAP endocytosis associated 2 | -0.26 | 3.333E-03 |
| ENSGACT00000021451 | si:ch211-113e8.11 | si:ch211-113e8.11 | -0.45 | 3.414E-03 |
| ENSGACT00000000223 | TRIM14 (5 of 74) | tripartite motif containing 14 | -0.36 | 3.472E-03 |
| ENSGACT00000015931 | CYP2W1 (5 of 5) | cytochrome P450 family 2 subfamily W polypeptide 1 | -0.44 | 3.482E-03 |
| ENSGACT00000021295 | B3GALT2 (2 of 2) | UDP-Gal:betaGlcNAc beta 13-galactosyltransferase polypeptide 2 | -0.66 | 3.521E-03 |
| ENSGACT00000020189 | trmt10c | tRNA methyltransferase 10 homolog C (S. cerevisiae) | -0.31 | 3.657E-03 |
| ENSGACT00000009880 | try (1 of 2) | trypsin | -0.32 | 3.681E-03 |
| ENSGACT00000013407 | pbx4 | pre-B-cell leukemia transcription factor 4 | -0.31 | 3.706E-03 |
| ENSGACT00000019112 | crygs3 | crystallin gamma S3 | -0.49 | 3.736E-03 |
| ENSGACT00000019922 | ap1s2 (2 of 2) | adaptor-related protein complex 1 sigma 2 subunit | -0.34 | 3.800E-03 |
| ENSGACT00000026352 | csrp2bp | csrp2 binding protein | -0.75 | 3.804E-03 |
| ENSGACT00000027496 | zw10 | zw10 kinetochore protein | -0.31 | 3.814E-03 |
| ENSGACT00000021697 | fbxo22 | F-box protein 22 | -0.65 | 3.880E-03 |
| ENSGACT00000018480 | C16orf70 | chromosome 16 open reading frame 70 | -0.53 | 3.887E-03 |
| ENSGACT00000015755 | tbc1d10aa | TBC1 domain family member 10Aa | -0.32 | 3.955E-03 |
| ENSGACT00000003070 | ldhd | lactate dehydrogenase D | -0.88 | 3.958E-03 |
| ENSGACT00000004185 | plekho1b | pleckstrin homology domain containing family O member 1b | -0.25 | 4.052E-03 |
| ENSGACT00000022149 | LEPROTL1 | leptin receptor overlapping transcript-like 1 | -0.47 | 4.066E-03 |
| ENSGACT00000015301 | MFSD3 | major facilitator superfamily domain containing 3 | -0.59 | 4.076E-03 |
| ENSGACT00000004129 | gpr12 | G protein-coupled receptor 12 | -0.59 | 4.171E-03 |
| ENSGACT00000000309 | tsn | translin | -0.30 | 4.252E-03 |
| ENSGACT00000006518 | pacsin1b | protein kinase C and casein kinase substrate in neurons 1b | -0.24 | 4.260E-03 |
| ENSGACT00000014825 | ptgr2 | prostaglandin reductase 2 | -0.36 | 4.306E-03 |
| ENSGACT00000011889 | slc26a10 | solute carrier family 26 member 10 | -0.38 | 4.341E-03 |
| ENSGACT00000010153 | loxl4 | lysyl oxidase-like 4 | -0.36 | 4.544E-03 |
| ENSGACT00000020822 | fbxo36b | F-box protein 36b | -0.25 | 4.566E-03 |
| ENSGACT00000000471 | foxp4 | forkhead box P4 | -0.43 | 4.624E-03 |
| ENSGACT00000025825 | LURAP1L | leucine rich adaptor protein 1-like | -0.65 | 4.689E-03 |
| ENSGACT00000009034 | plce1 | phospholipase C epsilon 1 | -0.51 | 4.721E-03 |
| ENSGACT00000012353 | mfsd5 | major facilitator superfamily domain containing 5 | -0.49 | 4.725E-03 |
| ENSGACT00000011081 | inpp5jb | inositol polyphosphate-5-phosphatase Jb | -0.31 | 4.731E-03 |
| ENSGACT00000007279 | fign | fidgetin | -0.34 | 4.766E-03 |
| ENSGACT00000015274 | s1pr3a | sphingosine-1-phosphate receptor 3a | -0.40 | 4.976E-03 |

Table S5. Genes that were significantly differentially expressed in both infected and SSRI-treated fish compared to controls. Four genes were found using p-values with a cut-off of 0.005 to identify significantly differentially expressed genes in each comparison. Gene ID, Gene name and Annotation refer to the identity of the transcript according to the *Gasterosteus aculeatus* BROAD-S1 genome assembly, genebuild release #83, NCBI accession ID: GCA_000180675.1. Log-fold change in expression in infected fish compared to controls or exposed fish compared to controls (logFC) and p-values (p-value) are presented. Genes differentially expressed compared to controls only in these two comparisons are in bold.

|  |  |  | **INFECTED vs CONTROLS** | | **SSRI-TREATED vs CONTROLS** | |
| --- | --- | --- | --- | --- | --- | --- |
| **Gene ID** | **Gene Name** | **Annotation** | **logFC** | **p-value** | **logFC** | **p-value** |
| **ENSGACT00000011412** | **mast3b** | **microtubule associated serine/threonine kinase 3b** | **0.33** | **5,020E-04** | **0.29** | **2,790E-04** |
| **ENSGACT00000026975** | **CCHCR1** | **coiled-coil alpha-helical rod protein 1** | **0.38** | **1,840E-03** | **0.37** | **4,240E-04** |
| ENSGACT00000008144 | ghdc | GH3 domain containing | 0.41 | 0.160e-02 | 0.71 | 1.720e-03 |
| ENSGACT00000012825 | slc38a10 | solute carrier family 38 member 10 | 0.59 | 0.167e-02 | 0.53 | 8.600e-04 |

Table S6. Genes differently expressed in the brain of SSRI-treated sticklebacks compared to controls. SSRI-treated individuals had 43 up-regulated and 9 down-regulated genes using p-values with a cut-off of 0.005 to identify significantly differentially expressed genes. Gene ID, Gene name and Annotation refer to the identity of the transcript according to the *Gasterosteus aculeatus* BROAD-S1 genome assembly, genebuild release #83, NCBI accession ID: GCA_000180675.1. Log-fold change in expression in SSRI-treated fish compared to controls (logFC) and p-values (p-value) are presented. Genes forming the genomic signature of SSRI-treated fish (genes differentially expressed only in this specific comparison) are in bold.

| **Gene ID** | **Gene Name** | **Annotation** | **logFC** | **p-value** |
| --- | --- | --- | --- | --- |
| **UP-REGULATED** | | | | |
| ENSGACT00000008144 | ghdc | GH3 domain containing | 0.72 | 1.718E-07 |
| ENSGACT00000014398 | ppp2r5ea | protein phosphatase 2 regulatory subunit B' epsilon isoform a | 0.61 | 1.905E-06 |
| ENSGACT00000004215 | RAB11FIP2 | RAB11 family interacting protein 2 (class I) | 1.15 | 4.008E-06 |
| **ENSGACT00000013336** | **zgc:92107** | **zgc:92107** | **0.94** | **7.182E-06** |
| ENSGACT00000012407 | prep | prolyl endopeptidase | 0.66 | 1.921E-05 |
| **ENSGACT00000015711** | **klhl12** | **kelch-like family member 12** | **0.55** | **4.458E-05** |
| ENSGACT00000026125 | fam161a | family with sequence similarity 161 member A | 0.67 | 6.011E-05 |
| ENSGACT00000007059 | slc5a8l | solute carrier family 5 (iodide transporter) member 8-like | 0.64 | 7.544E-05 |
| ENSGACT00000014388 | rtn4rl1b | reticulon 4 receptor-like 1b | 0.42 | 1.163E-04 |
| ENSGACT00000014419 | baiap2a | BAI1-associated protein 2a | 0.42 | 1.238E-04 |
| ENSGACT00000006518 | pacsin1b | protein kinase C and casein kinase substrate in neurons 1b | 0.32 | 2.243E-04 |
| ENSGACT00000027522 | PLEKHB1 | pleckstrin homology domain containing family B (evectins) member 1 | 0.81 | 2.479E-04 |
| ENSGACT00000011412 | mast3b | microtubule associated serine/threonine kinase 3b | 0.29 | 2.792E-04 |
| **ENSGACT00000012204** | **zgc:91944** | **zgc:91944** | **0.54** | **2.864E-04** |
| **ENSGACT00000003188** | **PAX2 (1 of 2)** | **paired box 2** | **0.34** | **3.134E-04** |
| ENSGACT00000026975 | CCHCR1 | coiled-coil alpha-helical rod protein 1 | 0.37 | 4.239E-04 |
| ENSGACT00000002153 | smg7 | SMG7 nonsense mediated mRNA decay factor | 0.57 | 4.322E-04 |
| ENSGACT00000022395 | myo9b | myosin IXb | 0.39 | 6.202E-04 |
| **ENSGACT00000006576** | **serpina1** | **serpin peptidase inhibitor clade A (alpha-1 antiproteinase antitrypsin) member 1** | **0.56** | **6.859E-04** |
| ENSGACT00000012825 | slc38a10 | solute carrier family 38 member 10 | 0.53 | 8.605E-04 |
| **ENSGACT00000014374** | **ankrd52a** | **ankyrin repeat domain 52a** | **0.54** | **9.444E-04** |
| **ENSGACT00000017487** | **snrpg** | **small nuclear ribonucleoprotein polypeptide G** | **0.29** | **9.461E-04** |
| **ENSGACT00000020942** | **tex9** | **testis expressed 9** | **0.55** | **9.863E-04** |
| ENSGACT00000009525 | tax1bp1a | Tax1 (human T-cell leukemia virus type I) binding protein 1a | 0.65 | 1.106E-03 |
| **ENSGACT00000013326** | **sulf2a** | **sulfatase 2a** | **0.97** | **1.196E-03** |
| ENSGACT00000014467 | osbpl1a | oxysterol binding protein-like 1A | 0.28 | 1.199E-03 |
| ENSGACT00000021551 | nip7 | NIP7 nucleolar pre-rRNA processing protein | 0.60 | 1.851E-03 |
| ENSGACT00000007714 | CASKIN1 (1 of 2) | CASK interacting protein 1 | 0.24 | 2.109E-03 |
| **ENSGACT00000004642** | **RGS22** | **regulator of G-protein signaling 22** | **0.46** | **2.414E-03** |
| **ENSGACT00000019804** | **DPP10 (3 of 3)** | **dipeptidyl-peptidase 10 (non-functional)** | **0.66** | **2.448E-03** |
| **ENSGACT00000014751** | **mrpl23** | **mitochondrial ribosomal protein L23** | **0.72** | **2.578E-03** |
| **ENSGACT00000001059** | **map4k2** | **mitogen-activated protein kinase kinase kinase kinase 2** | **0.67** | **2.625E-03** |
| **ENSGACT00000025595** | **lygl1** | **lysozyme g-like 1** | **0.33** | **2.634E-03** |
| **ENSGACT00000020860** | **saga** | **S-antigen, retina and pineal gland (arrestin) a** | **0.21** | **2.871E-03** |
| **ENSGACT00000007038** | **rarab** | **retinoic acid receptor alpha b** | **0.34** | **2.937E-03** |
| **ENSGACT00000005981** | **cks1b** | **CDC28 protein kinase regulatory subunit 1B** | **0.42** | **3.075E-03** |
| **ENSGACT00000023732** | **ghrb** | **growth hormone receptor b** | **0.41** | **3.372E-03** |
| **ENSGACT00000004688** | **olfml2a** | **olfactomedin-like 2A** | **0.64** | **3.582E-03** |
| **ENSGACT00000006216** | **si:ch211-114l13.9 (13 of 22)** | **si:ch211-114l13.9** | **0.55** | **3.677E-03** |
| **ENSGACT00000003203** | **ccr9a** | **chemokine (C-C motif) receptor 9a** | **0.66** | **3.948E-03** |
| **ENSGACT00000005507** | **adam8b** | **ADAM metallopeptidase domain 8b** | **0.58** | **4.606E-03** |
| **ENSGACT00000005372** | **slc39a4** | **solute carrier family 39 (zinc transporter) member 4** | **0.37** | **4.741E-03** |
| **ENSGACT00000010513** | **tmem237a** | **transmembrane protein 237a** | **0.34** | **4.811E-03** |
| **DOWN-REGULATED** | | | | |
| **ENSGACT00000023687** | **TRPC7** | **transient receptor potential cation channel subfamily C member 7** | **-0.64** | **3.246E-06** |
| **ENSGACT00000003040** | **lsm10** | **LSM10 U7 small nuclear RNA associated** | **-0.40** | **6.715E-04** |
| **ENSGACT00000003049** | **ndr2** | **nodal-related 2** | **-0.40** | **9.934E-04** |
| **ENSGACT00000027167** | **nf2b** | **neurofibromin 2b (merlin)** | **-0.25** | **1.152E-03** |
| **ENSGACT00000020887** | **tia1** | **TIA1 cytotoxic granule-associated RNA binding protein** | **-0.26** | **2.329E-03** |
| **ENSGACT00000001085** | **pygmb** | **phosphorylase glycogen muscle b** | **-0.34** | **2.555E-03** |
| **ENSGACT00000016674** | **wfikkn2a** | **info WAP follistatin/kazal immunoglobulin kunitz and netrin domain containing 2a** | **-0.26** | **3.117E-03** |
| **ENSGACT00000025355** | **plxnc1** | **plexin C1** | **-0.36** | **4.221E-03** |
| **ENSGACT00000010135** | **tefb** | **thyrotrophic embryonic factor b** | **-0.43** | **4.852E-03** |
